# Tradeoffs in optimal control capture patterns of human sensorimotor control and adaptation

**DOI:** 10.1101/730713

**Authors:** Tyler Cluff, Frederic Crevecoeur, Stephen H. Scott

## Abstract

Modern control theory highlights strategies that consider a range of factors, such as errors caused by environmental disturbances or inaccurate estimates of body or environmental dynamics. Here we reveal similar diversity in how humans naturally adapt and control their arm movements. We divided participants into groups based on how well they adapted to interaction loads during a single session of reaching movements. This classification revealed differences in how participants controlled their movements and responded to mechanical perturbations. Interestingly, variation in behaviour across good and partial adapters resembled simulations from stochastic and robust optimal feedback control, respectively, where the latter minimizes the effect of disturbances, including those introduced by inaccurate internal models of movement dynamics. In a second experiment, we varied the interaction loads over short time periods making it difficult to adapt. Under these conditions, participants who otherwise adapted well altered their behaviour and more closely resembled those using a robust control strategy. Taken together, the results suggest the diversity of how humans control and adapt their arm movements may reflect the accuracy of (or confidence in) their internal models. Our findings may open novel perspectives for interpreting motor behaviour in uncertain environments, or when neurologic dysfunction compromises motor adaptation.

## Introduction

Humans can perform a range of motor skills, from playing piano to throwing a baseball, but vary in how well they learn to perform these skills. Although it is widely accepted by coaches and music instructors that motor learning patterns vary between individuals^1–5^, studies typically focus on how factors, such as the structure or timing of practice, influence average patterns of adaptation across groups of participants^6–8^. This approach has had implications for how motor skills are taught^1^ and rehabilitated after neurologic injuries, disorders and diseases^9^.

Studies are only beginning to reveal the diversity of sensorimotor adaptation. It is becoming clear that healthy individuals differ in the rate they adapt their reaching movements when exposed to novel mechanical loads or visual disturbances within a single experimental session^10–15^. Interestingly, individuals who make the largest errors when first exposed to novel force disturbances tend to learn faster^12^. The authors argued that larger movement errors may accelerate adaptation by providing a better teaching signal. However, it is unclear why initial movement errors differ between individuals when exposed to the same visual or mechanical disturbances.

Healthy individuals also differ in the amount they adapt their reaching movements when they encounter novel mechanical loads. Although participants attain approximately 80% adaptation, on average, within minutes to hours of being exposed to novel movement dynamics, some participants display near-complete adaptation and others only partially adapt their movements with the same amount of practice^16–19^. Simulation studies suggest the nervous system is less sensitive to noise that corrupts motor commands or sensory receptors than it is to errors in its internal models of movement dynamics^20^. Indeed, even small internal model errors can produce unstable movements when moving in the presence of novel dynamics or external perturbations^21^. An emerging idea is that the nervous system may increase sensitivity to visual or somatosensory feedback to deal with incomplete learning^22, 23^, but it is unclear why and under what circumstances this strategy is beneficial for performing reaching movements.

Towards filling this gap, we demonstrate that principles of optimal feedback control capture tradeoffs in how people reach and respond to novel movement dynamics. Across two experiments, we demonstrate that patterns of human sensorimotor control and adaptation, measured in a single session of reaching movements, broadly resemble stochastic optimal feedback control and robust optimal feedback control. These two variants of optimal control differ in terms of how the controllers handle errors and uncertainty in their internal models of the body and environment. Taken together, the results and simulations illustrate how the accuracy of (or confidence in) internal models of the body and environment may influence the control strategies individuals select to interact with their environment.

## Results

### Experiment 1. Learning to move in the presence of novel interaction loads

We examined how healthy participants (N = 40; right handed) adapted their reaching movements when exposed to novel interaction loads. Participants first completed unloaded movements (Baseline) to three targets presented in random order (T1-T3, radius = 1 cm; Fig.1A). We positioned the targets so that the arm displacements required to reach them were identical across participants. Target 1 required shoulder motion (T1), Target 2 required combined shoulder and elbow motion (T2), and Target 3 required elbow motion (T3). We then introduced novel shoulder-joint loads (Fig.1 A-B; Adaptation) that were proportional to the angular velocity of the elbow joint (i.e., interaction loads)^24^. The interaction loads systematically disturbed the participant’s arm while they reached the training targets (T2-T3). Participants were instructed to reach the targets within 450-650 ms. Movement durations were determined as the time between when the participant’s hand left the start position (radius = 0.625 cm) to the time it entered the end target. The end target turned green to indicate successful timing, and blue (too slow) or red (too fast) if participants did not meet the timing demands.

**Figure 1.**
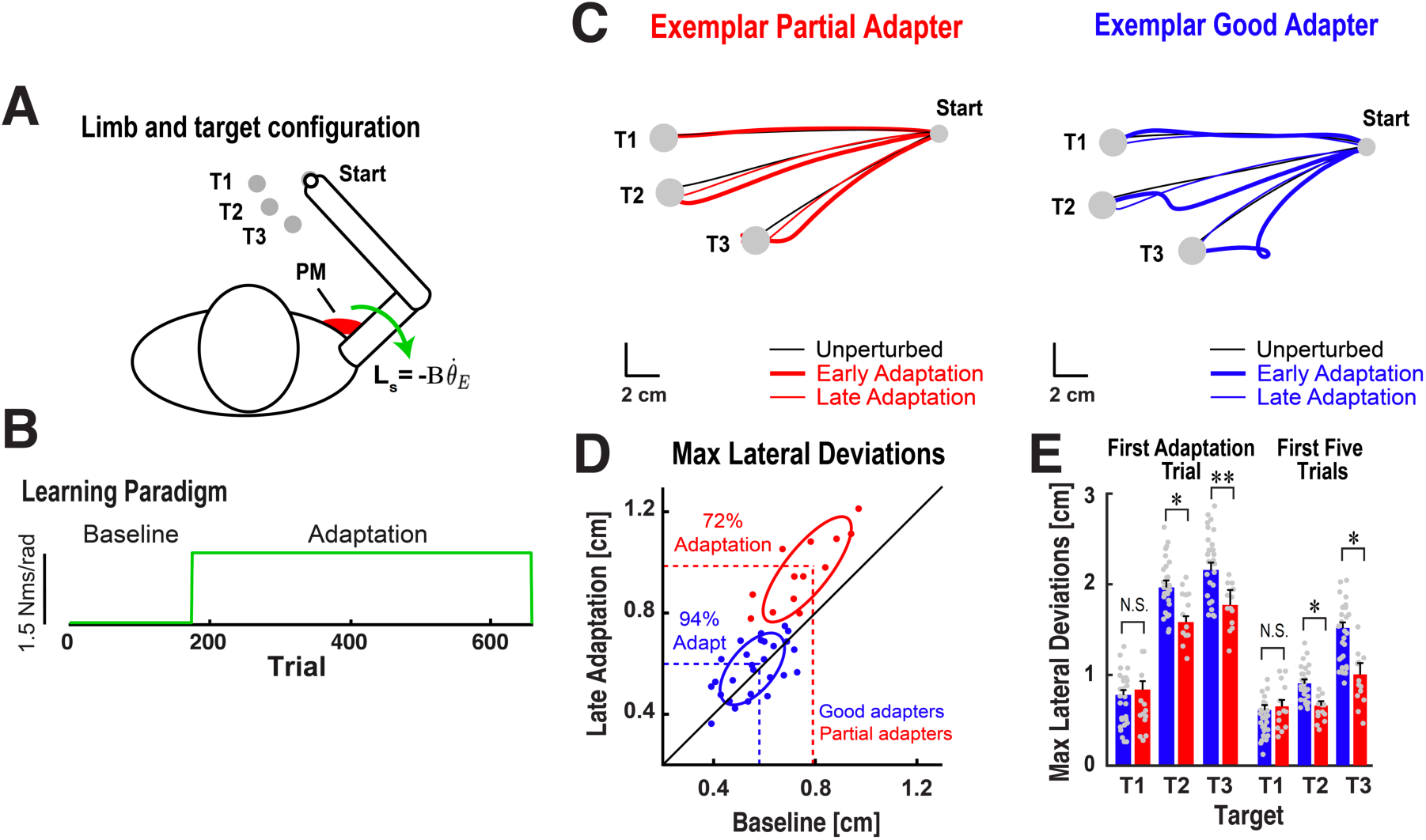
Behavioural task and adaptation paradigm. ***A***. Participants reached to two training targets (T2, T3) and a separate test target (T1). Green arrow represents novel shoulder loads (L_s_) applied in the adaptation block. ***B***. Time course of experiment. ***C*.** Hand paths for exemplar good and partial adapters. ***D***. Average hand deviations at targets T1-T3 during baseline trials and last 25 adaptation trials. ***E***. Average hand deviations from initial adaptation trials. \**p* < 0.05.

Participants also reached to a separate test target (T1) that required only shoulder flexion motion. We probed feedback responses by randomly disturbing the arm with step-torque perturbations that were applied the instant the participant’s hand left the start position (20% of trials at T1). The step-torque perturbations produced elbow joint extension and required a rapid correction to reach the target within the time constraints (-2Nm at shoulder and elbow; 10 ms sigmoidal ramp-to-constant profile)^25^. The same time constraints and performance feedback were used for unperturbed and perturbed reaching trials.

Elbow motion while reaching to T1 was limited (< 2**°**) and produced small, variable shoulder-joint loads during the adaptation phase (integrated shoulder load at T1 < 3% of loads at T2-T3). In principle, changes in muscle activity were not required to reach T1 when moving in the presence of the interaction loads. This design allowed us to probe feedback responses while minimizing the automatic gain scaling of muscle stretch responses^26–29^ caused by increases in muscle activity when countering the interaction loads to reach the training targets (T2-T3). In short, the design allowed us to test whether the amount participants compensated for the interaction loads while reaching the training targets (T2-T3) influenced how they responded to step-torque perturbations while reaching the test target (T1).

We first separated participants into two groups based on how well they compensated for the novel interaction loads. This metric captures systematic deviations in the hand paths of individual participants that reflect the accuracy of their internal models of arm and environmental dynamics^8, 16, 23, 30–34^. We classified participants as ‘good adapters’ if their peak hand-path deviations returned to near baseline levels by the end of training (Fig.1 C-D, single-sided *t*-test performed on individual participant data, *p* > 0.05). Participants were classified as ‘partial adapters’ if their hand path deviations remained larger than baseline levels (*p* < 0.05 for individual participants). As a group, good adapters displayed near complete adaptation (n = 27; 94.7 ± 3.9%), and partial adapters displayed incomplete adaptation (n = 13; 72.2 ± 4.9%, Supplementary Fig. 1).

Consistent with more accurate internal models of the novel interaction loads, good adapters displayed a larger increase in the agonist burst of their pectoralis major muscle (PM, shoulder flexor) while reaching target 3 (T3; Supplementary Fig. 2). The relationship between EMG bursts at the shoulder and elbow is well characterized for single-joint elbow movements^35–37^. Under natural conditions, elbow flexion motion produces an extension torque at the shoulder. When planning single-joint elbow movements, the nervous system counters extensor interaction torques by producing anticipatory activity of the shoulder flexor muscles. The interaction dynamics in our experiment accentuated the shoulder extensor torques produced during elbow flexion. Given that brachioradialis EMG (elbow flexor) was relatively constant during adaptation, the larger average increase in pectoralis major EMG displayed by good adapters suggests they exploited an internal model of the interaction dynamics.

Interestingly, we found that good adapters, who displayed near-complete adaptation to novel shoulder loads, also displayed larger peak hand-path deviations than partial adapters when the novel loads were first introduced (Fig. 1E, T2: *t*(38) = 2.18, *p* = 0.018, T3: *t*(38) = 2.45, *p* = 0.0096). Similar results were observed when broadening the analysis to the first 5 loaded movements to each training target (T2: *t*(38) = 1.88, *p* = 0.034; T3: *t*(38) = 1.81, *p* = 0.040). Despite some differences in peak shoulder loads, the overall mechanical demands of the task were qualitatively similar between good and partial adapters (Supplementary Fig. 3). Inertial properties of the arm were also similar across groups of good and partial adapters.

Collectively, our data highlight a link between movement errors when first exposed to novel interaction loads and the extent to which adaptive processes compensate for these loads. Does this link reflect differences in the way participants control and adapt their reaching movements? Optimal control theory may provide a framework to understand the link between initial and final performance, as there are many factors for the nervous system to consider when moving in the presence of novel dynamics. An influential theory for interpreting the function of the sensorimotor system is stochastic optimal feedback control (known as Linear-Quadratic-Gaussian, or extended LQG- control^38^). Stochastic optimal control uses a control policy to generate motor commands that represent the best way to perform an action (e.g., most accurate with minimal effort), assuming the controller’s internal model of arm and environmental dynamics is accurate (i.e., unbiased) and noise in the motor system follows a known distribution (Gaussian noise in the context of LQG). Even small biases in the internal model can cause movement errors when a stochastic optimal controller encounters loads that alter the dynamics of the arm or environment^39^.

Another class of optimal control models explicitly accounts for potential errors in the internal model when finding a solution for a motor task: robust optimal feedback control, which we formulate here in the framework of *H*_∞_-control^40, 41^. The principle in robust control is to set the feedback gains so that the control policy is as insensitive to disturbances as possible, including disturbances caused by biases or uncertainty in the internal model of body and environmental dynamics (i.e., the biological plant). Though insensitive to disturbances, robust control increases the cost of movement because the controller responds to all movement disturbances, regardless of how small or transient^41^. A stochastic optimal feedback controller ignores these deviations following the assumption it makes about the random (i.e., stochastic) nature of the disturbances.

In short, the control signals produced by stochastic optimal feedback control are less affected by noisy motor commands because the controller assumes noise disturbances are zero on average. The stochastic optimal feedback controller is also coupled with a minimum variance estimator (Kalman filter) that filters out noise in motor commands and sensory feedback when estimating the state of the arm. This allows the controller to perform efficient movements at the cost of making larger errors when the dynamics of the body or environment change unexpectedly. Robust control instead views any unmodeled disturbance as a perturbation, which results in control that is less efficient but also less sensitive to changes in the dynamics of the body or environment. In essence, stochastic and robust optimal feedback control lie at the extremes of a continuum of control strategies that differ in terms of how efficient versus sensitive the controller is to errors in its internal models of body and environmental dynamics.

Good and partial adapters were exposed to the same novel interaction loads but displayed differences in their average adaptation patterns. Within the context of stochastic and robust optimal feedback control, the observation that good adapters made larger initial errors but on average attained better final performance suggests they relied on their internal models of movement dynamics. This strategy would only be viable if they were capable of updating their internal model when confronted with interaction loads that altered the dynamics of the arm and environment. As a group, partial adapters appeared to compensate for the incomplete adaptation of their internal models by using a control strategy that was more robust to (potential) movement disturbances. By definition, robust control is less sensitive to internal model errors. This strategy would allow partial adapters to make smaller errors when the interaction loads were first introduced, as well as mitigate the impact of internal model errors throughout the (incomplete) adaptation process. Taken together, our data highlight a potential link between the amount participants adapt and the robustness of the strategy they select to control their reaching movements.

We formalized this link using a simplified model of the arm reaching to the same targets with the same time and accuracy demands as our human participants (Online Methods). The arm was controlled by stochastic and robust optimal feedback controllers paired with state estimators that helped circumvent the effect of noisy motor commands and noisy, time-delayed sensory feedback. Noise and cost parameters were taken from the literature or selected to generate motion patterns that qualitatively resembled human behaviour (see Table 1).

**Table 1:** Numerical values of parameters and their justification. Remark: the value of σ^2^ is equal to the elements of the noise covariance matrix of the motor noise. Thus the feedback noise has the same variance as the non-zero components of the motor noise, and the sensory covariance matrix has full rank in agreement with the assumptions of the Kalman filter^110^.

| Parameter | Value [unit] | Justification |
| --- | --- | --- |
| Inertia (arm; forearm and hand) | 0.14, 0.21 [kg m<sup>2</sup>] | Measured |
| Viscous friction | 0.14 [Nm s<sup>-1</sup>] | Reference<sup>39</sup> |
| Muscle time constant | 60 [ms] | Reference<sup>107</sup> |
| Motor noise covariance ($\Sigma_\xi$) | $0.025 \times BB^T$ | Qualitative match |
| Sensory noise covariance ($\Sigma_\zeta$) | $\sigma^2 I_{n \times n}$ | Reference<sup>38</sup> |
| Feedback delay | 50 [ms] | Latency of multi-joint FB |

#### Unperturbed Baseline Movements

Owing to its increased control gains, the robust controller produced baseline movements with faster and earlier peak hand velocities than the stochastic optimal controller (Fig. 2A). The robust optimal controller also generated more variable hand motion profiles than the stochastic optimal controller (Fig. 2A). Recall the two control models use the same cost function, time to complete the movement, and state space representation of the arm but solve the control problem in different ways. Increased variability emerges from the robust controller’s sensitivity to motor noise (process noise). Unlike stochastic optimal control, the robust controller does not assume disturbances are zero on average. Thus, the robust controller does not filter out disturbances arising from noise and treats any disturbance as a perturbation.

**Figure 2.**
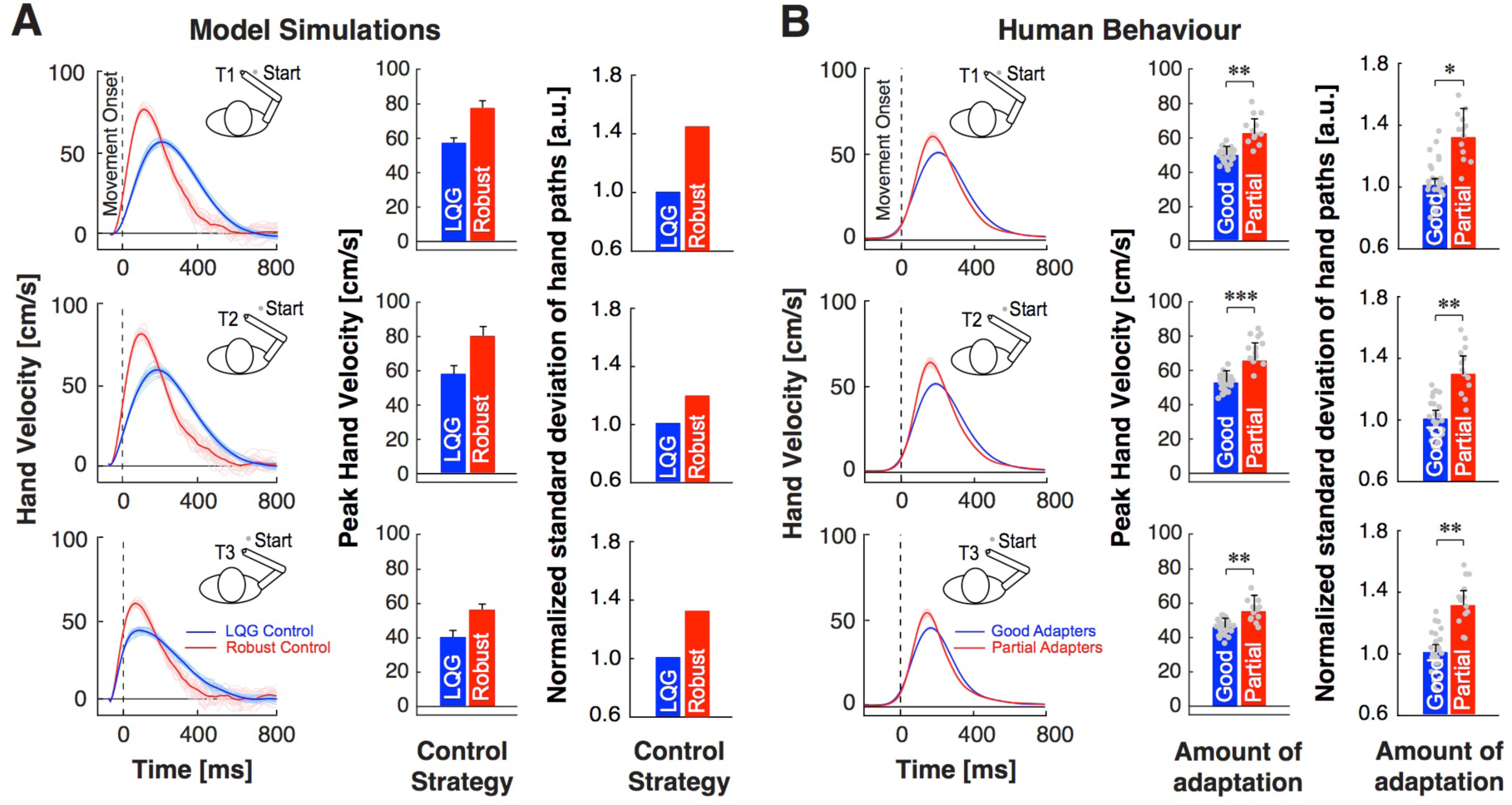
Baseline hand motion patterns. ***A***. Left: Simulated hand velocity profiles during baseline movements. Thin lines are individual trials; thick lines are average profiles. Right: Peak hand velocities and hand position variability. Data are normalized to variability of the stochastic optimal feedback controller (LQG). ***B*.** Left: Human hand velocity profiles (mean±SEM). Right: Peak hand velocities and position variability (mean±SEM). Data are normalized to the average variability displayed by good adapters. Dots represent individual participant averages. \**p* < 0.05, \*\**p* < 0.01, \*\*\**p* < 0.001.

Interestingly, we found the movements of robust and stochastic optimal controllers were paralleled by the behaviour of partial and good adapters. Partial adapters displayed faster peak hand velocities (Fig.2B, T1-T3: all *t*(38) > 3.05; all *p* < 0.01) and more variable movements than good adapters in the baseline phase of the experiment (T1-T3: all *t*(38) > 1.98, all *p* < 0.05).

#### Responses to Step-Torque Perturbations During Baseline Movements

Stochastic and robust optimal feedback controllers make fundamentally different assumptions about disturbances that arise during movement. Thus, we applied step-torque perturbations to view how the controllers responded to unexpected disturbances (Fig. 3A). The robust controller generated rapid and large corrective responses (Fig. 3C) to counter the perturbation and reach the target (Fig. 3B). When disturbed by the same step torque, the stochastic optimal controller generated larger control responses than the robust controller but later in movement (Fig. 3C). As a result, it displayed larger hand-path deviations and subsequent goal-directed corrections (Fig. 3B).

**Figure 3.**
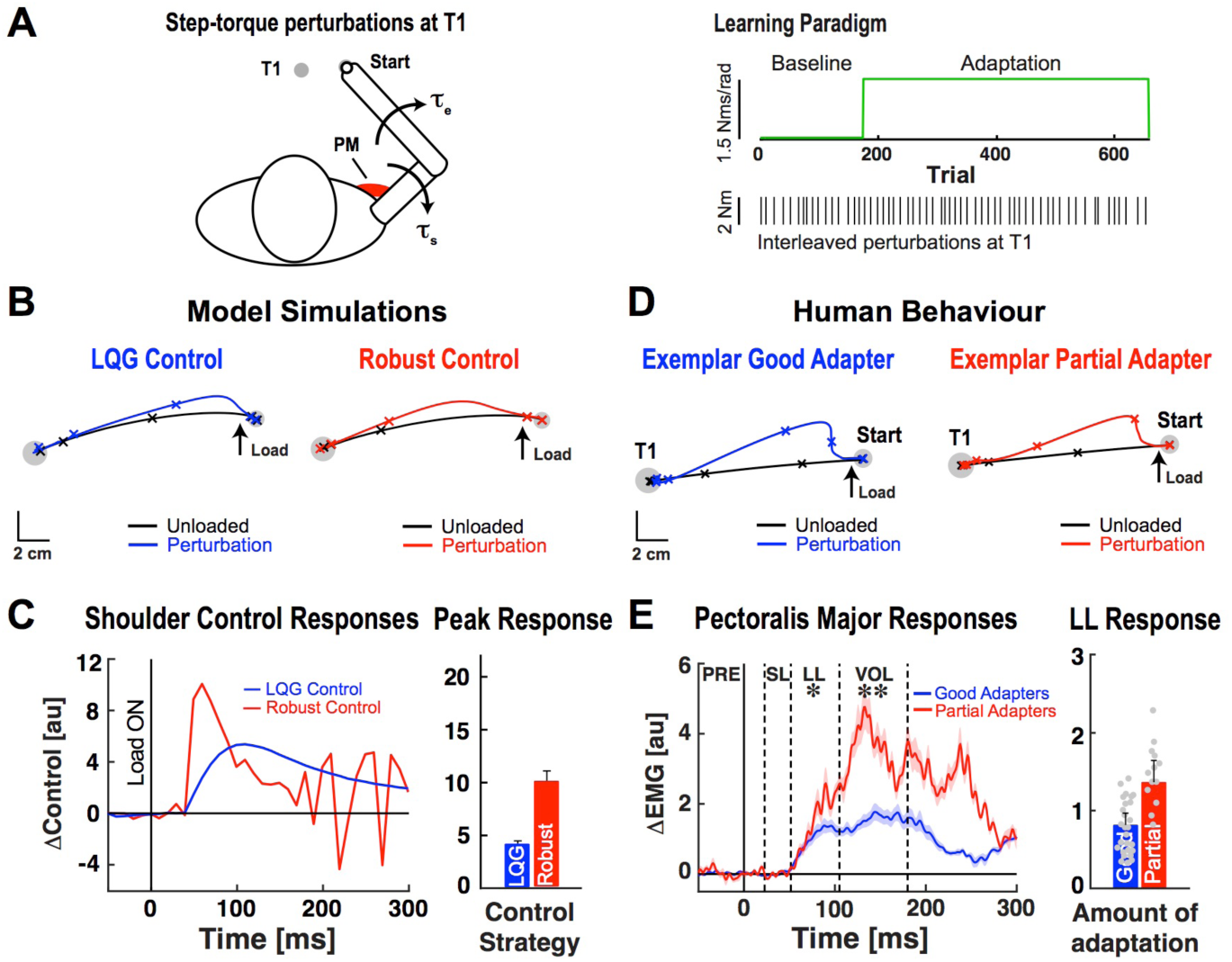
Baseline feedback corrections. ***A***. Left: Schematic of behavioural task and applied step-torque perturbations. Right: Time course of experiment. Black lines show interleaved step-torque perturbations applied while reaching the test target (T1). ***B***. Simulated hand paths during step-torque perturbation trials. ***C***. Shoulder control responses. Data are aligned to perturbation onset (Load ON). Bar graph shows the average peak in initial control response. ***D***. Hand motion patterns obtained from exemplar good and partial adapters during step-torque perturbation trials. Tick marks show the position of the hand every 200ms. ***E***. Pectoralis major stretch responses. Dashed vertical lines separate time phases of the muscle stretch response. Dots represent long-latency responses of individual participants. *\*p <* 0.05, \*\**p* < 0.01.

The behaviour of partial and good adapters paralleled how the robust and stochastic optimal controllers responded to step-torque disturbances, respectively. When their arm was disturbed by the step torque, partial adapters displayed smaller average peak hand deviations than good adapters (Fig. 3D, *t*(38) = 2.45, *p* < 0.01), and larger pectoralis major (PM) activity in the long-latency and voluntary time windows (Fig. 3E, LL = 50-105ms post-perturbation, *t*(38) = 2.05, *p* < 0.05; VOL = 105-180ms, *t*(38) = 2.22, *p* < 0.01). Note that differences in how the robust and stochastic optimal controllers responded to the step-torque perturbation emerged earlier (50-100 ms) than in the muscle responses of good and partial adapters (∼75-150 ms). Although our simplified model of muscle contraction (low pass filtered version of the control command) produces faster behavioural corrections, the EMG and behavioural results are consistent with the rise time of neural signals and evidence that it takes >100ms for humans to reach peak muscle activity when instructed to move as quickly as possible^42^. Muscle stretch responses were broadly consistent with model control signals at longer time scales (300+ ms; Supplementary Fig. 4).

### Adaptation to Novel Interaction Loads

#### Unperturbed Voluntary Movements

Next, we return to the issue of adaptation to novel interaction loads, as displayed in Figure 1C-D for human participants. Both controllers deviated from their baseline movements when moving with inaccurate knowledge of the novel interaction loads. However, the robust controller displayed smaller deviations than the stochastic optimal controller, just as partial adapters displayed smaller average hand path deviations than good adapters on the first adaptation trial (Fig.1C-D). We manipulated the accuracy of each controller’s internal model to examine how sensitive it was to imperfect knowledge of the interaction dynamics. The robust controller always displayed smaller peak hand path deviations than the stochastic optimal controller when moving with an inaccurate internal model of arm and environmental dynamics (Supplementary Fig. 5).

#### Feedback Corrections in the Presence of Novel Interaction Dynamics

We next explored how the controllers responded to step-torque perturbations when first exposed to novel interaction loads. When applied during baseline movements to T1, the step-torque extended the elbow while minimizing shoulder motion^43^. Combined elbow and shoulder flexor torques were required to counter the perturbation and reach the target. Recall the interaction loads produce shoulder loads proportional to the elbow’s angular velocity (elbow flexion causes shoulder extensor load). Elbow motion while reaching to T1 was limited (<2**°**) and produced small loads during reaching (<3% of load at T2-T3). When the same step torque was applied during adaptation, larger shoulder flexor torques were required to compensate for the extensor loads produced when countering the perturbation and flexing the elbow to reach the target. This approach allowed us to probe each controller’s sensitivity to disturbances and reliance on its internal model.

Introducing the novel load produced systematic errors in the internal models used by robust and stochastic optimal controllers. Internal model errors caused unexpected shoulder flexion motion when the step torque extended the elbow while moving in the presence of novel interaction loads. This caused both controllers to reduce their shoulder flexor response (Supplementary Fig. 6). Partial and good adapters also reduced their shoulder flexor responses when disturbed by the step-torque perturbation during initial adaptation trials (Supplementary Fig. 6).

We then compared model performance with human behaviour at the end of the adaptation block. For comparison, we constrained knowledge of the interaction load in our model simulations to the amount of adaptation displayed by our sample of good and partial adapters (Supplementary Fig. 7). Recall that good adapters, who better updated their internal models to compensate for interaction loads, displayed a larger average increase in shoulder flexor activity when producing pure elbow movements (T3). The stochastic optimal feedback controller also leveraged its adapted internal model. Before the onset of movement, it compensated for the expected interaction loads by producing a larger increase in shoulder flexor torque (relative and absolute) than the robust controller. This allowed the stochastic optimal controller to minimize movement disturbances while reaching target 3 (T3) in the presence of novel interaction loads (Supplementary Fig. 8). In short, the behaviour and control commands generated by robust and stochastic optimal controllers were broadly consistent with the behaviour of partial and good adapters. Differences in the adapted internal models of the stochastic and robust optimal feedback controllers influenced how they responded to step-torque disturbances. Due to its sensitivity to motion disturbances and partially adapted internal model, the robust controller decreased its shoulder flexor response when the step-torque extended the elbow and produced unexpected shoulder flexion motion. In contrast, the stochastic optimal controller’s internal model was more accurate and it generated a larger shoulder flexor response to account for extensor torques produced when countering the step-torque and flexing the elbow to reach the target. As a result, it was less disturbed by the perturbation than the robust controller (Fig.4A-B). In short, the robust controller produced control signals that were more sensitive to individual patterns of joint motion when moving in the presence of novel interaction dynamics. Stochastic optimal control generated responses that were sensitive to the estimated inter-joint dynamics.

**Figure 4.**
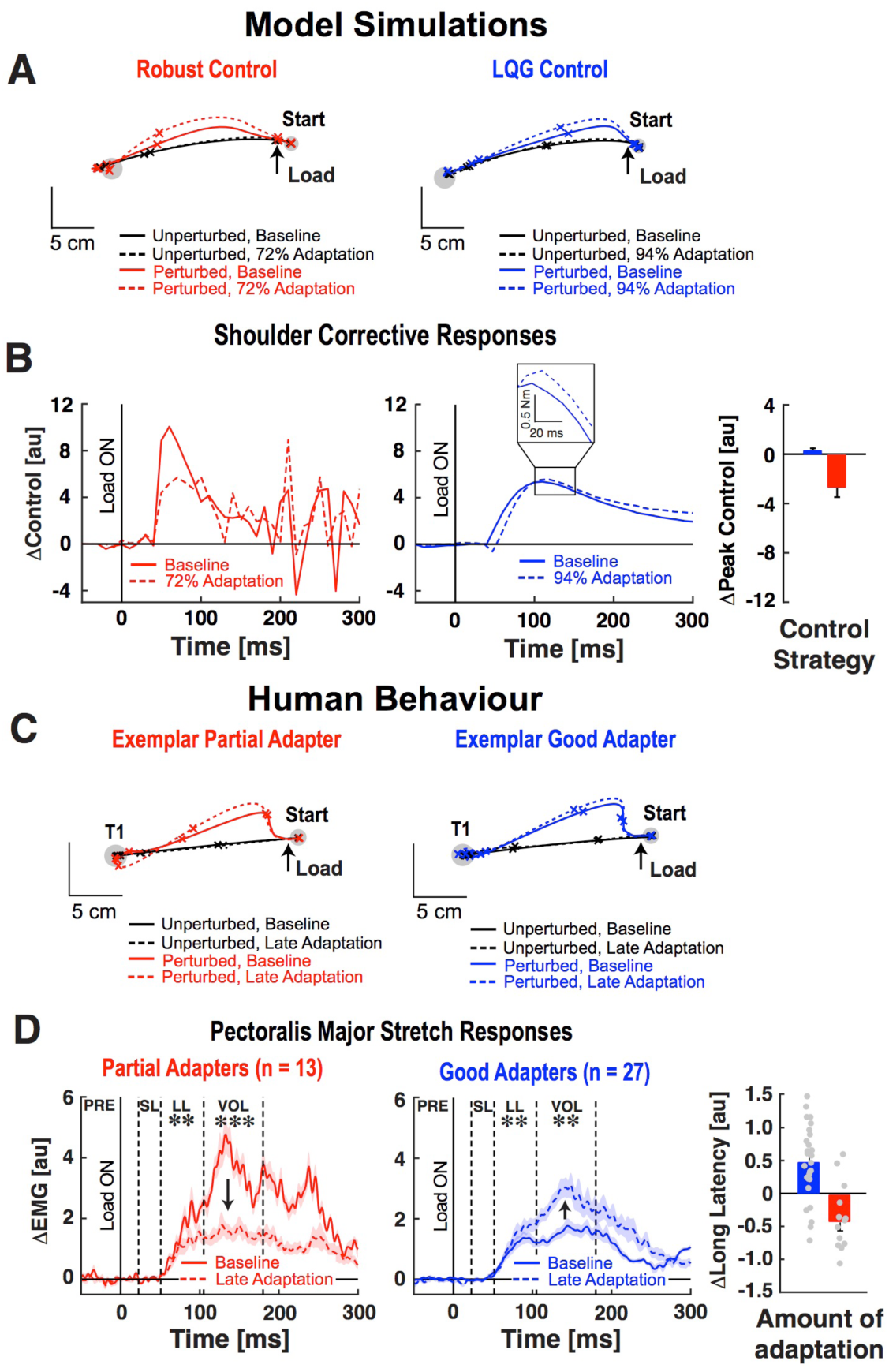
Comparison of baseline and adapted feedback responses. ***A***. Simulated hand paths during step-torque perturbation trials. ***B***. Shoulder control responses. Inset shows the LQG-control response. Bar graph shows average change in peak control response. ***C***. Hand motion obtained from exemplar good and partial adapters during step-torque perturbation trials. Tick marks show hand positions every 200ms. ***D***. Pectoralis major stretch responses. Dashed vertical lines separate time phases of the muscle stretch response. Dots represent the change in long-latency response of individual participants. \*\**p* < 0.01, \*\*\**p* < 0.001.

The model responses were paralleled by how good and partial adapters responded to step-torque perturbations at the end of the adaptation phase. On average, good adapters increased their PM responses (Fig. 4D, LL & VOL time windows, *t*(26) > 2.73, *p* < 0.01) to counter shoulder extensor loads produced when correcting elbow motion. In contrast, partial adapters still reduced their PM responses at the end of the adaptation phase (LL & VOL, *t*(12) > 2.26, *p* < 0.01; elbow responses in Supplementary Fig. 9). Good adapters displayed adapted hand paths that were disturbed less by the step-torque perturbations than those displayed by partial adapters (Fig.4C, *t*(38) = 2.86, *p* < 0.01).

### Experiment 2. Dealing with Errors Caused by Abrupt Changes in Dynamics

The results of Experiment 1 show that, on average, participants who compensated less for novel interaction loads seemed to rely on feedback about individual patterns of joint motion rather than generating responses suitable for the dynamics when reaching and responding to step-torque disturbances. Previous studies have reported a decrease in learning rates and increased reliance on sensory feedback when moving in unpredictable dynamical environments^22, 44^.

Motivated by these results, we performed a second experiment to test whether participants would make their strategy more robust when exposed to interaction dynamics that changed frequently. The rapid changes in dynamics would induce frequent internal model errors and make it challenging to produce accurate movements when relying on a stochastic optimal feedback control strategy. This control strategy is sensitive to changes in dynamics because it does not account for errors or uncertainty in internal models. Thus, if good adapters use such a strategy, we expected that frequent and abrupt changes in dynamics would have a larger impact on their behaviour compared to that of partial adapters.

#### Adaptation to constant interaction dynamics

Participants were initially exposed to the same interaction load as Experiment 1 (Fig. 5A). Our first step was to determine whether they behaved and adapted in similar ways as participants in Experiment 1. Figure 5B shows representative hand paths from participants who displayed partial and near-complete compensation for the interaction loads. Taking the same approach as Experiment 1, we separated participants into groups based on how well they compensated for the interaction loads. This analysis reproduced the results of Experiment 1. Good adapters (Fig. 5C, n = 18; 92.3 ± 6.9%) better compensated for the interaction loads than partial adapters (n = 12; 76.1 ± 5.5%, Supplementary Fig. 10). Consistent with Experiment 1, good adapters also made larger errors when the interaction loads were first introduced (Fig. 5D, First trial at T2-T3: all *t*(28) > 1.88, *p* < 0.05, First 5 trials at T2-T3: all *t*(28) > 2.71, *p* < 0.05).

**Figure 5.**
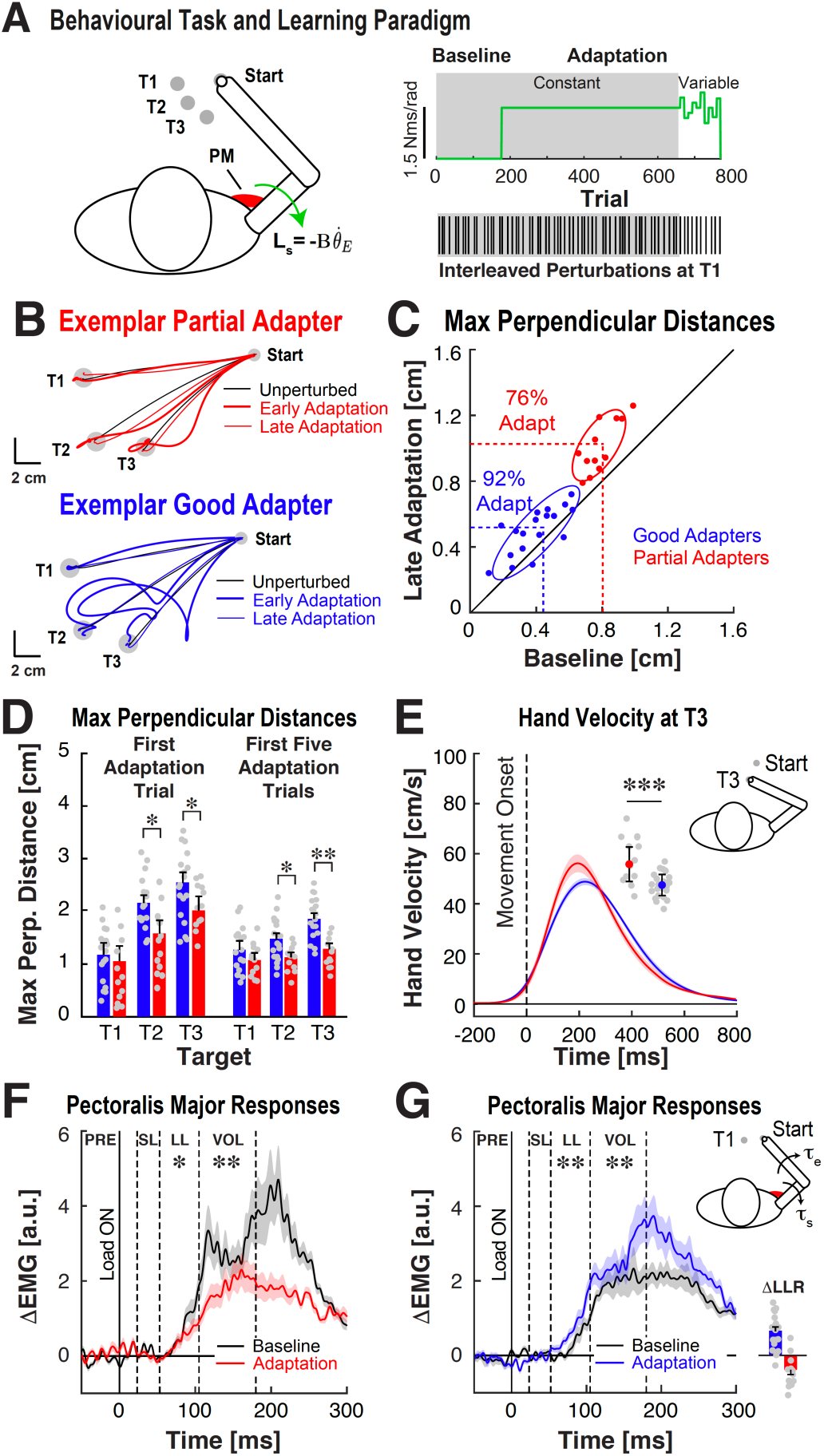
Behaviour and feedback corrections in Experiment 2. ***A***. Behavioural task and time course of experiment. Participants reached to two training targets (T2, T3) and a separate test target (T1). Green arrow represents novel shoulder loads (L_s_) applied in the adaptation block. **B**. Hand paths for exemplar partial and good adapters. ***C***. Average hand deviations at targets T1-T3 during baseline trials and last 25 adaptation trials at each target. ***D***. Average hand deviations from initial adaptation trials. ***E***. Average velocity profiles observed for partial and good adapters. Error bars represent the SEM. Blue and red dots represent average peak velocity for groups of good and partial adapters (± 1 SEM). Grey dots represent individual subject averages. ***F*.** Average pectoralis major stretch responses displayed by partial adapters. Dashed vertical lines separate time phases of the muscle stretch response. ***G***. Average pectoralis major stretch responses displayed by good adapters. Data plotted in the same format as ***F***. Bar graphs represent average change in LLR between baseline and late adaptation. Dots represent individual participant data. \**p* < 0.05, \*\**p* < 0.01.

We also examined the baseline hand velocity profiles of good and partial adapters. Recall that in Experiment 1, partial adapters displayed faster, earlier peaks in their hand velocity profiles than good adapters. Consistent with Experiment 1, partial adapters again displayed faster and earlier peaks in their hand velocity profiles than good adapters (Fig. 5E, T3: *t*(28) = 4.77; *p* < 0.001).

We probed the feedback responses of good and partial adapters by measuring the amplitude of their muscle responses when disturbed by the same step-torque perturbations used in Experiment 1 (20% of trials at T1). Partial adapters again displayed larger baseline PM responses than good adapters when their arm was disturbed by a step-torque perturbation (LL & VOL, all t(28) > 2.20, *p* < 0.05). As in Experiment 1, good adapters leveraged their internal model when responding to the step torque during the adaptation phase. They increased their PM responses (Fig. 5F-G, LL & VOL, all *t*(17) > 2.89, *p* < 0.01) in anticipation of shoulder extensor loads produced when countering the step torque (i.e., flexing the elbow) to reach the target. Partial adapters reduced their PM responses during adaptation (LL & VOL, all *t*(11) < 2.36, *p* < 0.05).

The classification criterion separated participants into two groups by design. However, this was only for analysis purpose and it is clear that the range of behavioural traits is by no means dichotomous. To show this, we investigated the relationship between the amount participants in Experiments 1 (n = 40) and 2 (n = 30) adapted their hand paths (%adaptation) and modulated their PM stretch responses between the baseline and late phases of adaptation (ΔEMG). We found a significant correlation between the amount participants adapted their movements and changes in their PM responses (Fig. 6; R^2^ = 0.44, *p* < 0.001, intercept = -2.35, slope = 0.031, F_1,68_ = 53, *p* < 0.001).

**Figure 6.**
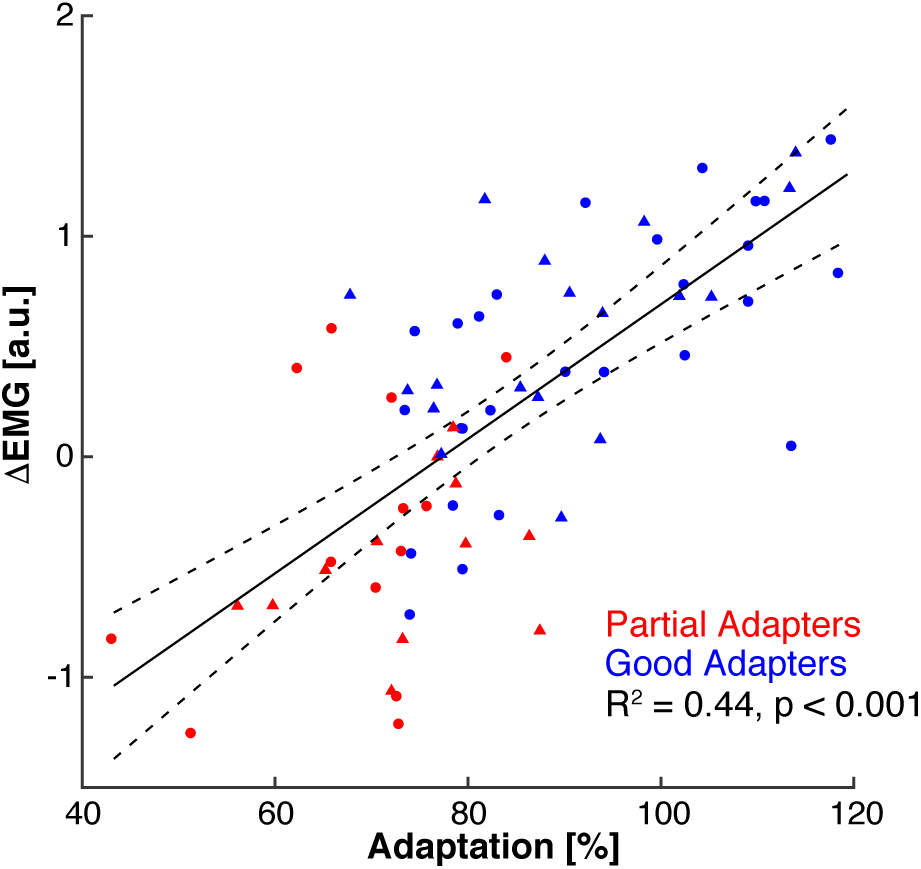
Relationship between adaptation (%) and modulation of pectoralis major stretch responses between the baseline and late adaptation phases of the experiment (ΔEMG). Circles represent participants from Experiment 1, triangles represent participants from Experiment 2.

#### Adapting to abrupt changes in interaction dynamics

After participants completed the adaptation phase, we varied the load coefficient after each block of 11 trials (Fig. 7A, 1.05-1.95 Nm.s/rad, mean load coefficient = 1.5 Nm.s/rad). We probed feedback responses with the same step-torque perturbations (-2 Nm, 10 ms sigmoidal ramp; 20% of trials at T1).

**Figure 7.**
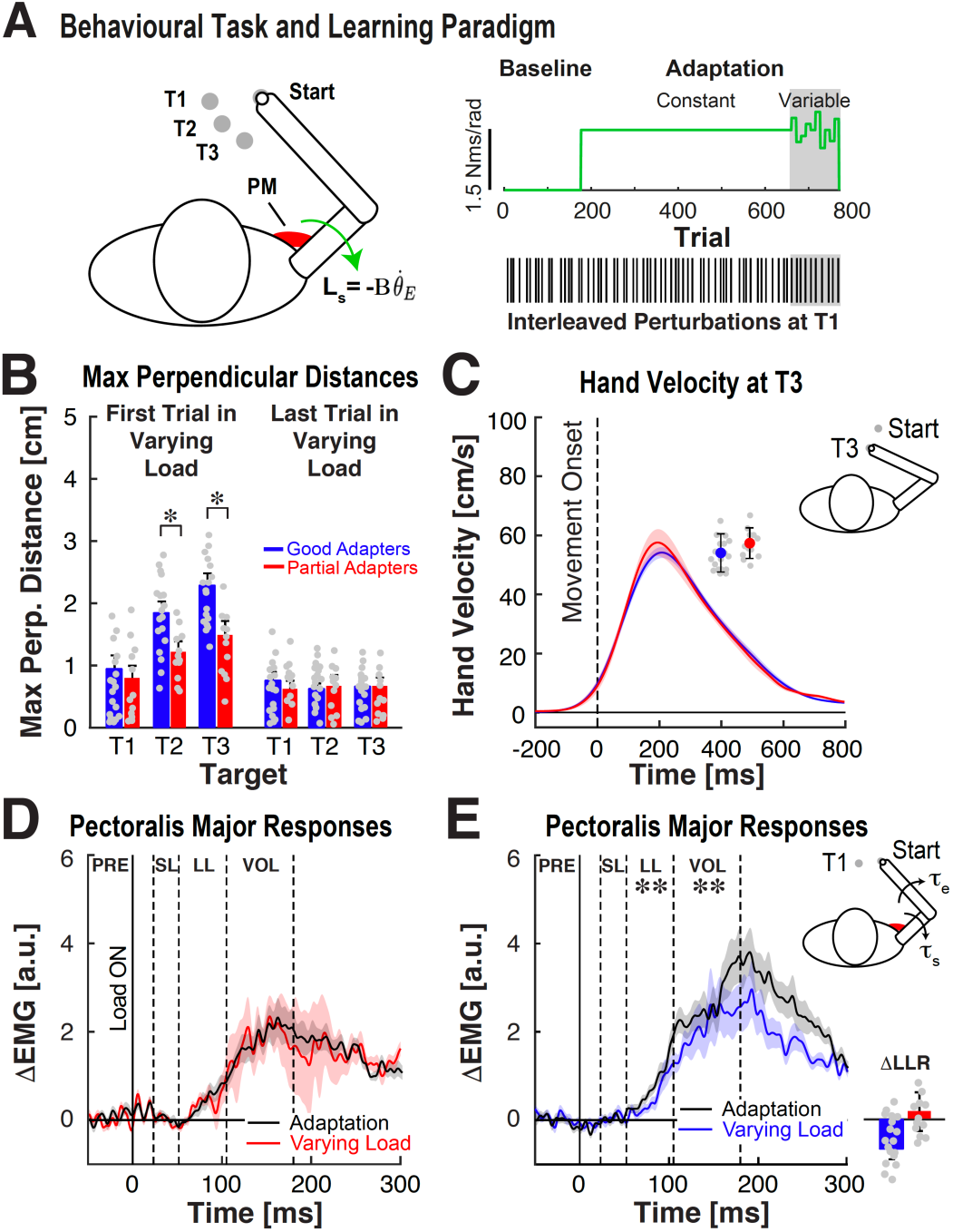
Behaviour and feedback corrections when exposed to frequent and abrupt changes in interaction loads. ***A***. Behavioural task and time course of experiment. ***B*.** Average hand path deviations displayed by good and partial adapters when exposed to interaction dynamics that changed abruptly over short blocks of trials. ***C***. Average hand velocity profiles observed for partial (n =12) and good adapters (n = 18). Error bars represent the SEM. Blue and red dots represent average peak velocity for groups of good and partial adapters. Grey dots represent individual subject averages. ***D*.** Average pectoralis major stretch responses displayed by partial adapters when exposed to constant and varying interaction dynamics. Dashed vertical lines separate time phases of the muscle stretch response. ***E***. Average pectoralis major stretch responses displayed by good adapters. Data plotted in same format as ***D***. Bar graphs represent average change in LLR. Dots represent individual participant data. \**p* < 0.05, \*\**p* < 0.01, \*\*\**p* < 0.001.

Consistent with our hypothesis, good adapters made larger errors when first exposed to new interaction dynamics (Fig. 7B, T2-T3, *t*(28) > 2.17, *p* < 0.05). However, their hand paths became similar to those displayed by partial adapters after prolonged exposure to changes in interaction dynamics (Fig. 7B, T2-T3, *t*(28) < 0.29, *p* > 0.49). Good adapters increased and shifted their peak hand velocity to earlier in movement. Thus, their hand velocity profile became more similar to the profile displayed by partial adapters (Fig. 7C, *t*(28) = 0.33, *p* = 0.74) and predicted by robust optimal feedback control.

When their arm was disturbed by the step-torque perturbation while reaching to target 1 (T1), partial adapters showed little change in their PM responses compared to when exposed to constant interaction dynamics (Fig. 7D, LL & VOL, all *t*(11) < 1.81, *p* > 0.18). In contrast, good adapters reduced their PM responses (Fig. 7E, LL & VOL, all *t*(17) > 3.84, *p* < 0.01). Good adapters also increased the amplitude of their brachioradialis responses (Supplementary Figure 11, LL: *t*(17) = 2.36, *p* = 0.029, VOL: *t*(17) = 1.86, *p* = 0.08), suggesting their control strategy became less dependent on multi-joint motion. Thus, participants seemed to use a robust-like strategy when dealing with an environment that changed frequently and caused internal model errors.

## Discussion

In agreement with a growing body of literature^for review see 45^, we found large variation in how healthy adults adapted their arm movements to counter novel interaction loads. In order to understand how this variation impacted the control of reaching movements, we classified participants as good or partial adapters based on how well they compensated for novel interaction loads. This approach captured several features of behaviour. As a group, partial adapters made smaller errors than good adapters when exposed to novel interaction loads. During baseline movements, partial adapters displayed faster peak hand velocities, as well as larger muscle responses when disturbed by step-torque perturbations. These variable patterns of sensorimotor control and adaptation were observed despite identical task instructions, performance feedback, similar mechanical properties of the arm and overall demands of the task.

The behaviour displayed by good and partial adapters was broadly consistent with stochastic and robust optimal feedback control, respectively. The control models had identical physical properties, noise parameters, feedback delays and cost functions, but make different assumptions about disturbances that arise during movement (inclusion or not of potential internal model bias or uncertainty). Collectively, our models and data highlight a tradeoff between the efficiency of the control strategy and how sensitive it is to disturbances, including those introduced by imperfect or uncertain internal models of movement dynamics.

We believe the diversity of control and adaptation patterns reflects the extent to which participants relied on model-based control of their reaching movements. Previous work has shown that relying on internal models can cause larger errors when the task changes unexpectedly^10^. Consistent with this argument, introducing novel interaction dynamics caused good adapters to make larger movement deviations than partial adapters, who seemed to mitigate internal model errors (or lack of confidence in their internal models) by relying more on feedback about the motion of their shoulder and elbow joints. Strategies play a role in how people learn and perform cognitive skills^46–49^ and make decisions^50, 51^. We highlight that strategies may also play a role in how people adapt and control their arm movements when briefly exposed to novel movement dynamics.

Our results reveal a link between the adaptation and control of movement dynamics. Variable patterns of adaptation have also been reported when healthy adults receive offline error feedback while adapting their reaching movements to a visuomotor rotation^52^. The study by Palidis and colleagues^60^ revealed a correlation between learning rates and the amplitude of error-related potentials recorded over parietal cortex. The authors argued that this correlation may stem from sensory prediction errors, such that individuals who adapted more rapidly to the visuomotor rotation relied on internal models to estimate the outcomes of their actions. Moreover, a recent study demonstrated that error-related muscle activity correlates with how much participants update their next movement when exposed to a force field^12^. Collectively, these findings suggest individuals who rely more on model-based control use error-related feedback to better adapt their reaching movements. Our second experiment examined whether participants would alter their control strategies when exposed to less predictable dynamics. We found that good adapters made larger errors than partial adapters when first exposed to unpredictable dynamics, but eventually attained similar performance. Moreover, good adapters increased their movement velocity and displayed patterns of behaviour that more closely resembled robust control. Similar changes in movement velocity have been reported when environmental forces vary unpredictably between trials^53^. When disturbed by a step torque, good adapters displayed increased control gains and shoulder responses that depended less on the imposed relationship between elbow and shoulder motion and more on individual patterns of joint motion.

Robust control is a costly but cautious strategy. The apparent change in strategy highlighted in Experiment 2 suggests good adapters were less confident in their estimate of the changing interaction dynamics. However, their shoulder responses were larger (and elbow responses smaller) than those displayed by partial adapters. This suggests participants made their strategy more robust but still leveraged knowledge of the interaction dynamics – though to a lesser extent. Taken together, our findings support the idea that people rely more on sensory feedback when interacting with less predictable movement dynamics^22, 54, 55^.

Our experiments examined how participants controlled and adapted their reaching movements while interacting with a robot in a single session of reaching movements. Thus, we may only provide a static snapshot of tradeoffs between learning and control. This approach is common in studies that examine how humans adapt their motor actions^56–58^, respond to mechanical disturbances^59–61^, and shape beliefs about whether errors arise from changes in properties of the body or environment^62^ within a brief period of practice. One exception is the work by Nezafat and colleagues^63^ who measured force-field adaptation over 4 weeks. Interestingly, the activity of the deep cerebellar nuclei changed throughout adaptation despite behaviour reaching a steady state within the first practice session. This raises the question of whether participants can improve their internal models and alter their control strategies (particularly partial adapters), or instead display stable patterns of learning and control with long term adaptation. It is also unclear whether similar tradeoffs apply in tasks that depend less on dynamics (e.g., playing piano) or are performed outside of the laboratory.

The ability to adapt internal models may not always transfer to other aspects of motor function. A recent study by Stark-Inbar and colleagues^64^ demonstrated the rate individuals adapt their movements when exposed to a visuomotor rotation does not correlate with how fast they learn motor sequences. Evidence suggests these tasks rely on different neural substrates and learning mechanisms^65^. Although it is well-established that visuomotor and force-field adaptation rely on error-based mechanisms, they impose different control problems for the nervous system. In principle, robust strategies can be developed for a broad class of disturbances. Dependent on the nature of internal models of cursor motion, participants may display similar variation in the control strategies they select when adapting to visuomotor rotations. Indeed, recent evidence suggests participants may upregulate their feedback responses when exposed to a visuomotor rotation^66^. Interestingly, slower learners displayed larger spinal reflexes when their arm was disturbed by a mechanical perturbation. The extent to which increased gains arise from uncertain or inaccurate internal models of cursor motion and to what extent control strategies can be altered by instruction are open questions for future work.

We argue that the behaviour of good and partial adapters align with features of feedback control strategies that make different assumptions about movement disturbances. Previous studies have argued the nervous system regulates the limb’s intrinsic resistance to disturbances (i.e., impedance) by modulating muscle co-activation in a way that depends on the direction and stability of novel force environments^34, 67–71^. Here we found that long-latency responses varied with the accuracy of an individual’s internal model of the interaction dynamics, which caused partial adapters to decrease their pectoralis major stretch responses due to unexpected shoulder motion. Importantly, this result is referenced to each participant’s baseline stretch responses, suggesting partial adapters used a strategy that was more robust (on average) and driven by increased feedback gains rather than limb impedance. Standard approaches also calculate the arm’s impedance by averaging motion over time periods as long as 300 ms after the onset of a perturbation^71^. Thus, it is difficult to attribute changes in force responses that involve neural feedback gains^72–76^ solely to changes in the arm’s intrinsic stiffness or damping. The current results revealed that partial adapters displayed higher levels of muscle activity. However, past work has shown that moderate to high levels of co-contraction cause only small changes in the arm’s impedance^39^. When disturbed by a perturbation, the authors reported differences in joint motion that emerged >80 ms after perturbation onset and correlated with the amplitude of neural feedback responses involving the spinal cord and brain. In short, our current results suggest the link between learning and control is reflected in the earliest time period that neural control policies contribute to corrective responses. We are pursuing whether and under what circumstances spinal reflexes may facilitate robust control strategies.

The match between our model simulations and human EMG was not perfect. Our model is a linearized version of the arm’s biomechanics driven by actuators that can produce flexor and extensor control torques. The control torques were a low-pass filtered version of the control signals obtained from stochastic and robust optimal feedback policies. The models did not consider the length- or velocity-dependent properties of muscle^77, 78^, nor did they consider the role of biarticular or antagonist muscles. Furthermore, we did not fit the models to participant data but set the parameters to examine the principle that uncertain or inaccurate internal models can influence the control strategies that participants select to interact with their environment. Despite being relatively simple, the models qualitatively captured features of human sensorimotor control and adaptation.

We used a performance criterion to categorize individuals as good or partial adapters. However, we really view this as a continuum where individuals may tradeoff the robustness and efficiency of movement. This continuum became clear when viewing how changes in muscle responses varied with the amount of adaptation. While some overlap between the behaviour of good and partial adapters may reflect that robust control is capable of similar performance with slightly lower levels of adaptation, some is also due to our simple performance criterion (i.e., statistical contrasts misclassified some participants). Nevertheless, this approach allowed us to test model predictions and capture the most pronounced traits of what is indeed a continuum of strategies.

The continuum of strategies reveals a link between the control and adaptation patterns of the dominant arm. It may also reveal an interesting parallel with work on inter-limb differences in reaching movements^79–85^. Past work suggests the dominant arm may (on average) be better suited for rapid learning of internal models of predictable dynamics, while the non-dominant arm may rely on non-specific control strategies that are less sensitive to unpredictable changes in dynamics. If the nervous system indeed uses model-based strategies to control the dominant arm, then we expect it may also display efficient corrective responses that more closely resemble stochastic optimal control. Conversely, the non-dominant arm may display upregulated feedback responses that more closely resemble robust control. Less clear is whether differences in control persist when the arms attain similar levels of performance in unpredictable dynamical environments^86^.

Our results may have implications for understanding impairments that accompany neurologic disorders. Several studies have highlighted the cerebellum’s role in internal models for motor control and learning^33, 87–90^. Cerebellar damage can impair internal models for controlling movements^91–95^ and reduce the rate and amount of adaptation when exposed to novel loads^31, 96–101^. An interesting question is whether individuals with cerebellar damage would also select a robust strategy when their internal models are inaccurate. There is also considerable variation in the recovery of sensory and motor function after stroke^102^. Simulations of the reaching movements of hemiparetic stroke survivors suggest impairment may arise from the inability to accurately represent properties of the arm^20^. Stroke survivors may select a robust strategy to downplay the effects of errors in the internal model of their stroke-affected limb—if sensory processing is left intact. However, when stroke impairs feedback and internal models, stochastic optimal and robust control strategies would both be substantially impaired, leading to deficits in the control and adaptation of arm movements.

Selecting a control strategy that is appropriate for the task and environment may be an important factor in motor performance. When movement conditions are highly predictable, such as a pitcher throwing a baseball, relying on an internal model may reduce the variability and energetic cost of movement. In contrast, robust feedback control may improve performance when dealing with unpredictable disturbances while walking down a crowded street or wrestling an opponent. At the extreme, it may be favourable to switch between strategies depending on the situation, similar to the change in behaviour displayed by good adapters when the interaction dynamics changed frequently and made it difficult to adapt. Robust control may improve performance if an opponent is close when shooting a basketball, whereas reducing neural feedback gains and leveraging internal models may be desirable for a free throw when there is little concern of being disturbed.

## Materials and Methods

### Participants

Forty healthy right-handed adults participated in Experiment 1 (17 males, 21-44 years old). The Queen’s University Research Ethics Committee approved the experimental procedures. Thirty healthy right-handed adults participated in Experiment 2 (20 males, 20-37 years old). The protocol for Experiment 2 was approved by the University of Calgary Conjoint Health and Research Ethics Board. Participants gave written informed consent before participating and received monetary compensation for their time. Experiments were completed in 90 minutes, including participant preparation time.

### Apparatus and Behavioural Task

Participants made reaching movements while seated with their right arm supported in a robotic exoskeleton that can apply selective mechanical loads to the shoulder and/or elbow joints (KINARM, BKIN Technologies)^24, 103^. Targets and hand-position feedback (white cursor, diameter = 0.8 cm) were projected into the plane of the participant’s arm using a semi-silvered mirror. Direct vision of the arm and hand were occluded throughout the experiment.

The behavioural task is shown in Figure 1*A*. Participants began the trial by moving their hand to the start target (0.625 cm radius) and briefly holding this position (1000 ± 500 ms, uniformly distributed). One of three potential targets was then shown in random order (T1-T3; 1 cm radius). We instructed participants to reach the target within 450-650ms. Post-trial feedback about movement time was provided during a 1s hold period at the spatial goal. The target turned green if the reach was completed on time and turned blue (too slow) or red (too fast) if the participant did not reach the target within this time window. Movement times were calculated as the time between when the participant’s hand left the start position and entered the spatial goal. An elbow extensor background load (-1 Nm; 500 ms sigmoidal ramp-up) was applied at the start of each trial. The background load ramped down after movement completion (500ms sigmoidal ramp-down) and remained off for 2 s between trials.

### Experiment 1. Learning to move in the presence of novel interaction loads

Participants first performed unloaded movements (Baseline: 176 trials). We then applied novel interaction loads at the shoulder (Adaptation: 484 trials) that were proportional to the angular velocity of the elbow. The shoulder loads were applied at each time sample (1 ms) according to the following equation:

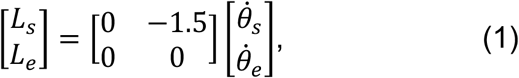

where [*L_s_ L_e_*]*^T^* represent interaction loads (Nm) applied by the robotic device, and 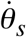 and 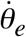 represent the angular velocities of the shoulder and elbow (rad/s). Note the shoulder loads were applied in the direction opposite the elbow’s angular velocity, such that elbow flexion produced shoulder extensor loads.

The experiment was divided into 15 blocks (∼4-5 min/block) separated into three phases: baseline (4 blocks), early adaptation (5 blocks), and late adaptation (6 blocks). Each block consisted of 44 trials, including 20 movements to T1, 12 movements to T2, and 12 movements to T3. Four of the 20 movements to T1 were interleaved step-torque perturbations. Two-minute rest breaks were provided after every 4 blocks. Additional rest breaks were provided at the participant’s request.

Our paradigm uses training targets that require combined shoulder and elbow motion (T2; 10° shoulder flexion, 10° elbow flexion), or only elbow motion (T3, 20° elbow flexion). Participants also reached to a separate test target that required only shoulder flexion motion (T1; 20° shoulder flexion). Elbow motion while reaching T1 was limited (<2**°**) and produced only small residual loads while participants reached to the test target (integrated shoulder load at T1 < 3% of integrated shoulder loads experienced at T2 and T3). This allowed us to maintain relatively similar amounts of muscle activity while participants reached T1. Our design allowed us to test if knowledge of the interaction loads expressed while reaching the training targets (T2, T3) was reflected in shoulder muscle responses when disturbed by a step-torque perturbation while reaching to the test target (T1).

### Feedback Responses to Mechanical Perturbations

Feedback responses were measured on random trials by applying step-torque perturbations that extended the elbow (-2 Nm at shoulder and elbow; 10ms sigmoidal rise-time) while participants reached the test target. The step-torque perturbations were triggered the instant the participant’s hand left the start position. Equal amounts of torque were applied at the shoulder and elbow joints^44^. Participants were instructed to reach the spatial goal within the same time window as unperturbed trials and received the same post-trial feedback.

### Experiment 2. Interacting with rapidly changing interaction dynamics

Experiment 2 used the same behavioural task as Experiment 1. Participants (n = 30) first performed unloaded movements. We then introduced the same interaction loads at the shoulder (load constant = -1.5 Nm.s/rad). After participants completed the adaptation phase, we varied the load coefficient in blocks of 11 trials. Each block included an interleaved step-torque perturbation that displaced the elbow into extension while participants reached to the test target (T1). The size of the load coefficient was altered in each block of trials while maintaining the same average coefficient as when exposed to constant interaction dynamics. Other aspects of the experiment were identical to Experiment 1.

### Data Analysis

#### Kinematic recordings and learning analysis

Shoulder and elbow joint kinematics were sampled at 1 kHz and digitally low-pass filtered (2^nd^-order, dual-pass Butterworth, 30 Hz effective cutoff). We calculated the perpendicular distance between the hand and a straight line joining the start position and goal at each time sample^104^. We measured changes in reaching patterns by comparing peak hand-path deviations in the baseline block with the last 25 reaching movements to each target in the adaptation block. Participants were classified as good adapters if their hand paths returned to near baseline levels of performance at the end of adaptation (single-sided paired t-test for individual participants, *p*>0.05), or partial adapters if their hand paths still deviated from their baseline movements at the end of adaptation (single-sided t-test for individual participants, *p*<0.05). The amount of learning (% adaptation) was determined by comparing hand-path deviations at the end of learning with baseline movements:

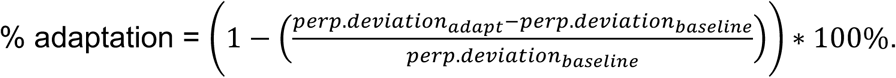

The hand velocity profile and peak velocities in the direction of the target were computed on a trial-by-trial basis for each participant. Averages were then compared across groups of good and partial adapters (single-sided unpaired t-test, *p*<0.05). We then normalized the hand-path variability displayed by partial adapters during movements to each separate target (T1-T3) to the average tangential hand-path variability displayed by good adapters while reaching the same target. We contrasted the normalized hand-path variability between good and partial adapters (single-sided unpaired t-test, *p*<0.05).

#### EMG Recordings and Analysis

EMG of the upper limb muscles was recorded using bipolar surface electrodes (DE 2.1 Single Differential Electrode, Delsys). EMG signals were amplified online (gain = 10^3^-10^4^) and sampled at 1-kHz. We recorded the activity of the monoarticular elbow muscles (brachioradialis, elbow flexor; triceps lateralis, elbow extensor) and shoulder muscles (pectoralis major, shoulder flexor; posterior deltoid, shoulder extensor). We also recorded from the biarticular flexor muscles (biceps brachii, short head) and extensor muscles (triceps longus). We cleaned the skin with alcohol before attaching electrodes over each muscle. A ground electrode was placed over the patella or elbow.

EMG data were aligned to perturbation onset, band-pass filtered (3^rd^-order, 20-450 Hz pass band, dual-pass Butterworth) and full-wave rectified. We normalized each muscle’s rectified activity during reaching to its activity while maintaining postural control at the start position against 1 Nm extensor (brachioradialis, biceps brachii, pectoralis major) or flexor background loads (triceps lateralis, triceps longus, posterior deltoid). The posture task was completed at the start of the experiment (five 15s trials for each load direction). Additional details about the EMG normalization can be found in Cluff and Scott (2013)^72^.

Muscle stretch responses were aligned to the onset of the step-torque perturbation. We subtracted the average EMG in unperturbed trials from perturbed trials (ΔEMG). We then calculated each muscle’s average activity (ΔEMG) in predefined time windows on a trial-by-trial basis for each participant^72, 105^: pre-perturbation (PRE: -50–0 ms), short-latency (SL: 25–50 ms), long-latency (LL: 50– 105 ms) and early voluntary (VOL: 105–180 ms). We examined how good and partial adapters altered their muscle responses by comparing average EMG responses to the step-torque perturbation in each time window between the baseline and late adaptation phases (paired t-tests). Similarly, we compared hand motion caused by the step-torque perturbation between baseline and late adaptation trials. EMG data were smoothed with a bidirectional 5-ms moving average filter for display purposes. All statistical tests were performed on unsmoothed EMG data.

### Optimal Feedback Control Models

The control models are based on a linear approximation of the arm moving in the horizontal plane. We calculated the inertia of the arm segments based on the physical characteristics of our human participants using standard anthropometric estimates^106^. We combined these inertial estimates with the inertial properties of the robotic exoskeleton. The mechanical model of the arm was coupled with a first-order, low-pass filter linking control input to muscle force to approximate the dynamics of muscular contraction (time constant: 60ms)^107^. We simulated reaching movements to the same targets (T1-T3), with the same time (450-650ms) and accuracy demands (target diameter = 2 cm) as our human experiments.

We consider two optimal control algorithms. The first follows the literature on sensorimotor control and learning, and is based on stochastic optimal control, or linear-quadratic-Gaussian regulator (LQG control)^108^. This control model assumes plant uncertainty is captured by a random variable that follows a known (Gaussian) distribution. The second control algorithm is based on robust control, formulated here in the context of *H*͚-optimal control. The robust control algorithm achieves optimal attenuation of unknown external disturbances, including uncertainty or errors in the internal model of the plant (i.e., the upper limb)^40^. This control model has recently been used to describe human sensorimotor control^109^ and can be designed to handle errors or uncertainty in the internal model of the arm, including the error that we induce experimentally by introducing the novel interaction load.

The discrete time control system is expressed in the following equation, in which *x_t_* represents the state vector including joint angles, velocities and torques:

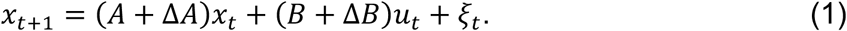

The matrices *A* and *B* are the modeled (or expected) dynamics, Δ*A* and Δ*B* are the model errors, and ξ_t_ is a zero-mean Gaussian disturbance with covariance Σ_ξ_]. The discrete time system is coupled with the feedback equation:

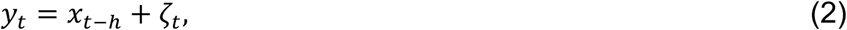

where *h* is the feedback delay expressed in number of time samples and ζ*_t_* is the additive Gaussian noise that corrupts sensory feedback with covariance Σ_ζ_. Temporal delays were handled with standard system augmentation^60^. Equation 1 can be transformed into:

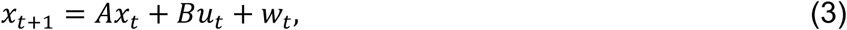

where the term *w_t_*: = Δ*Ax_t_* + Δ*Bu_t_* + *ξ_t_* captures the noise disturbances, as well as un-modeled limb dynamics. With these definitions, the control problem consists of minimizing the following cost-function^40^:

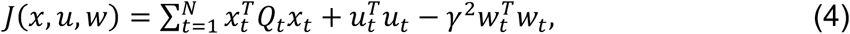

while using the incomplete state information from Equation 2.

Stochastic optimal control and robust control solve this problem in different ways. In the framework of stochastic optimal control, it is assumed that Δ*A* = Δ*B* = 0, and the control solution minimizes the expected value of *J*(*x*,*u*,*w*). A well-known solution is given by (extended) LQG control^38^. The robust control approach considers that Δ*A* and Δ*B* can be non-zero, and the controller minimizes the impact of this unknown disturbance. The control law is derived in the worst-case scenario formulated as a zero-sum dynamic game. The full derivation of the robust controller was provided in detail by Basar and Bernhard^40^. Observe that in our experiment, the internal model error following the introduction of the novel load applies to the state variables by mapping elbow velocity into shoulder torque. As a consequence, our experiment corresponds to the situation where Δ*B* = 0 and Δ*A* ≠ 0.

The two controllers are coupled with a state estimator, which is necessary to handle noise in sensory signals as well as temporal delays induced by the transmission and processing of neural signals (50ms delay, Equation 2). The optimal state estimator for the stochastic optimal controller is the standard Kalman filter^76^. The optimal estimator for the robust controller differs from the Kalman filter because the plant uncertainty also affects how feedback signals are used to correct the prediction about the present state of the limb. However, the general form of this estimator is similar to the Kalman filter in the sense that it is a linear combination of the previous state estimate, control input and feedback vector^40^.

The cost matrices (*Q_t_*) penalize all state variables to ensure the robust control problem is not ill-posed^3^. We applied a constant cost on velocity and muscle force (numerical values:30 ×(*velocity*)^2^ and 0.1 ×(*torque*)^2^), whereas the running cost on joint angles followed a quadratic build up towards the final cost (final cost: 6 × 10^3^ × (*position error*)^2^). These matrices were adjusted to ensure the robust control problem is well defined (i.e., the cost matrix *Q_t_* is invertible)^40^, as well as to generate movement trajectories that qualitatively resembled the behaviour of our experimental participants. The cost parameters were kept identical for the two control designs, and thus the changes in behavior indeed reflect differences in the control strategies. The parameter γ is jointly optimized with the control function and represents the optimal level of disturbance attenuation. The two controllers converge to a linear control law applied to the estimated state. Table 1 lists the model parameters and provides numerical values used in the simulations. Paralleling the experimental procedures, we first characterized reaching movements during a baseline period consisting of unloaded movements (Δ*A* = 0). Step-torque perturbations were simulated with abrupt changes in the external torque applied during the simulation runs^39, 60^.

We then examined how the models performed when reaching to the same targets in the presence of a novel interaction load (Δ*A* ≠ 0) that was identical to our human experiments and was proportional to the instantaneous angular velocity of the elbow. We simulated learning by manipulating knowledge of the interaction load. We varied knowledge of the parameter, *L* (-1.5 Nm.s/rad), by multiplying with a scalar (0, 0.25, 0.50, 0.75, 1.00), which is the constant that maps elbow angular velocities onto interaction loads applied at the shoulder (corresponding to 0, 10, 25, 50, 75, and 100% knowledge of the load coefficient). For comparison purposes, we constrained knowledge of the novel load to the amount of learning displayed by good (94% adaptation) and partial adapters (72% adaptation). We also looked at how the two control algorithms respond to the same step-torque perturbations before (25% learning) and after adapting to novel interaction loads (robust control: 72% learning; LQG control: 94% learning). Finally, we calculated tangential hand velocity profiles for the models, as well as hand-path variability during movement (t = 0-800 ms) for the LQG and robust control algorithms. We then expressed hand-path variability for the robust controller as a percentage of that noted for the LQG controller (i.e., 1 = identical hand-path variability for both controllers).

Note the structure of the cost function was partially constrained by the requirements of the game-theoretic approach used for the robust controller (i.e., *Q_t_* must be invertible). Importantly, this approach allowed us to use the same cost function structure for simulations with the robust and LQG controllers. Thus, the free parameters were the amount of noise corrupting the motor command (process noise) and sensory feedback (observation noise). We reduced the number of free parameters by assuming sensory and motor noise terms were equivalent.

Changing the cost function and noise parameters will alter the quantitative correspondence between model simulations and human behavior. Although human behavior may not be captured quantitatively by other parameter settings, the fundamental principle that robust control reduces sensitivity to internal model errors, and the trade-off between efficiency and robustness remains true across all parameter settings. Thus, the difference in how robust and LQG-control solve the control problem has an overarching impact on how the controllers respond to external perturbations and novel interaction loads.

## Author contributions

T.C. performed the experiments; T.C., F.C., S.H.S. analyzed the data, F.C. performed the simulations, T.C., F.C., S.HS. wrote the paper.

## Code and data availability

Custom computer code was used to control the experimental task, analyze behavioural data and perform the computational modeling. Computer code and behavioral data are available upon request to the corresponding author.

## Acknowledgements

We thank Kim Moore and Justin Peterson for their technical support. We thank members of the LIMB Lab for their helpful feedback and comments. We would also like to thank Julien Hendrickx for critical comments on an earlier version of this manuscript. This work was supported by NSERC Discovery grants to T.C. and S.H.S.

