## Supplemental Figures for "Tradeoffs in optimal control capture patterns of human sensorimotor control and adaptation"

### Supplementary Figures and Captions

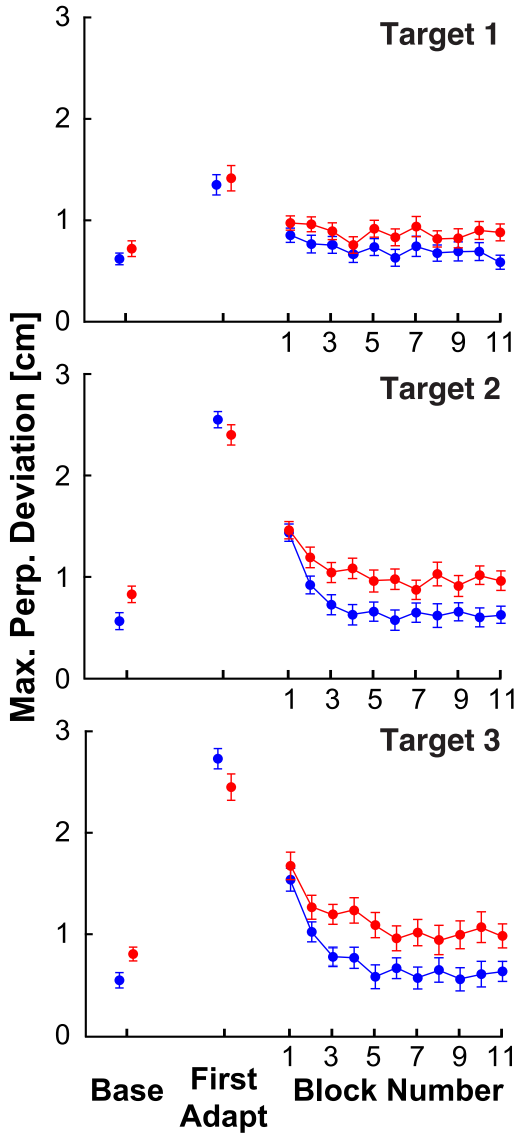


**Supplementary Fig. 1. Adaptation profiles in Experiment 1.** Maximum perpendicular deviations were calculated across late baseline, initial exposure to the interaction load at each target, and across each block of trials during the adaptation phase. Error bars represent ± 1 SEM.

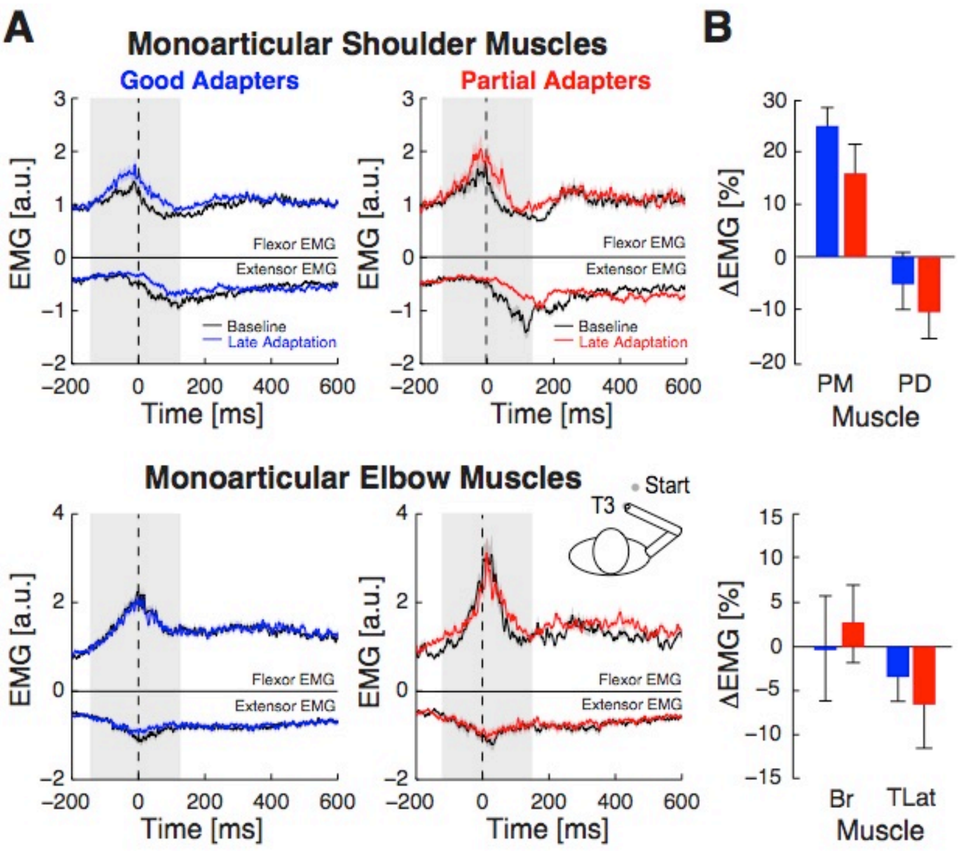


**Supplementary Fig. 2. Adaptation of muscle activity when reaching with novel interaction loads. A.** Average activity of shoulder (top row) and elbow muscles (bottom row) observed while groups of good and partial adapters reached to target 3. Black lines represent average muscle activity in the baseline phase. Coloured lines represent average muscle activity at the end of adaptation (last 25 trials at T3). Dashed vertical lines represent movement onset. Shaded grey region corresponds to time epoch used to integrate EMG during the agonist burst. **B**. ΔEMG signal (%) obtained by normalizing the difference in each muscle's agonist burst in the adaptation phase to its agonist burst in the baseline phase. Good adapters displayed a larger increase in pectoralis major EMG to counter shoulder extensor torques produced while flexing their elbow joint to reach target 3 (T3).

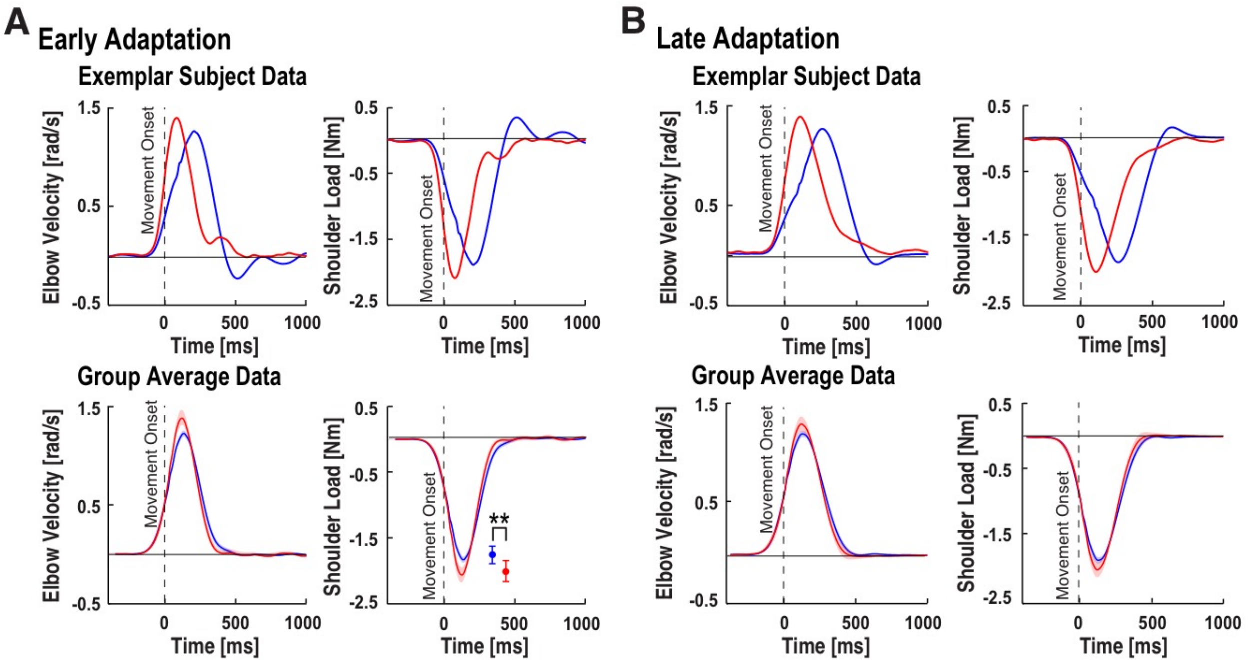


**Supplementary Fig. 3. Shoulder loads applied while reaching to target 3 (T3). A.** Early adaptation phase (first ten trials). Top row: Elbow flexion velocity (left column) and resulting shoulder extensor torques (right column) for a representative partial (red trace) and good adapter (blue trace). The partial adapter generated movements with faster and earlier peak elbow joint velocities than the exemplar good adapter. Integrated shoulder loads were similar across groups of good and partial adapters (T2: $t(38) = 1.52$, $p = 0.14$; T3: $t(38) = 0.47$, $p = 0.64$) despite differences in peak elbow velocities and shoulder extensor loads (T2: $t(38) = 3.43$, $p = 0.0020$, T3: $t(38) = 3.13$, $p = 0.0031$). Bottom row: Group averages for good adapters (n = 27) and partial adapters (n = 13). Data presented in the same format as above. **B.** Late adaptation phase (last 25 trials to T3). Data presented in the same format as **A.** We computed the inertia of upper limb segments based on the physical characteristics of our participants using standard anthropometric estimates[42]. We combined these estimates with the inertial properties of the robot. The inertia of the forearm and hand (unpaired single-sided t-test; $t(38) = 0.71$, $p = 0.49$), as well as upper arm segments ($t(38) = 0.71$, $p = 0.37$) were not significantly different between groups of good and partial adapters.

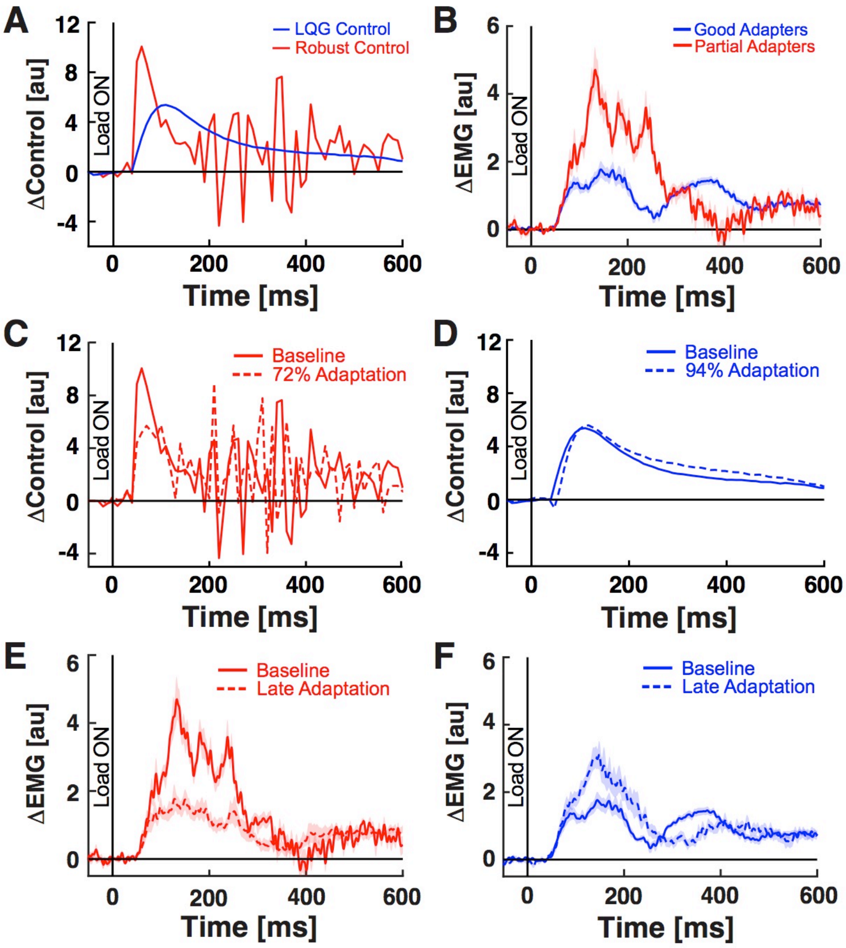


**Supplementary Fig. 4. Control signals generated by robust and stochastic optimal feedback controllers and human EMG. *A.*** Baseline control responses of robust and stochastic optimal feedback controllers when disturbed by a step-torque perturbation. ***B.*** Baseline EMG responses of partial and good adapters when disturbed by a step-torque perturbation. ***C-D.*** Comparison of baseline (solid lines) and adapted (dashed lines) responses of robust (***C***) and stochastic optimal feedback controllers (***D***) when disturbed by step-torque perturbations. ***E-F.*** Comparison of baseline (solid lines) and adapted (dashed lines) EMG responses of partial (***E***) and good adapters (***F***) when disturbed by step-torque perturbations.

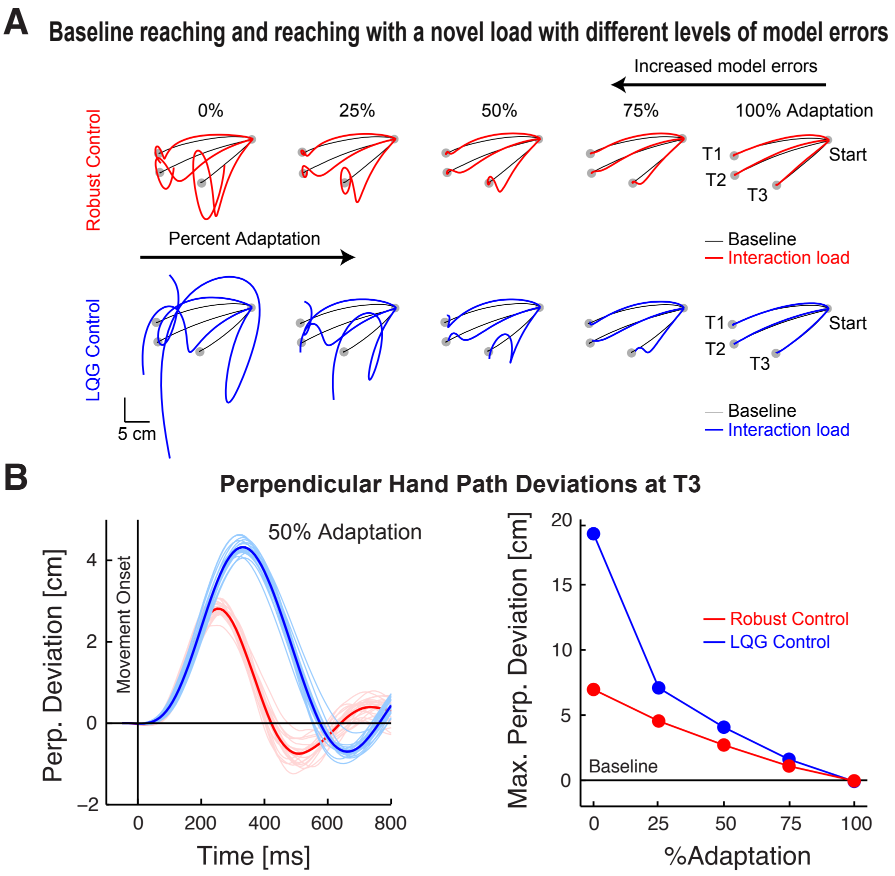


**Supplementary Fig. 5. Reaching movements generated by robust and stochastic (LQG) optimal feedback controllers. *A.*** Simulated hand paths during unloaded (baseline) and loaded (adaptation) reaching movements for robust and stochastic optimal feedback controllers (LQG). Thin black trace is the average hand path during baseline movements (n = 20 trials); thick coloured trace is the average hand path when reaching with novel interaction loads (n = 20 trials). The interaction load caused both controllers to deviate from their baseline movements (0% adaptation). Lateral deviations decreased systematically when the controllers had improved internal models of the interaction dynamics (0-100% adaptation). ***B.*** *Left panel*: Perpendicular hand path deviations while reaching to T3 with 50% adaptation to novel interaction loads. Thin lines represent individual trials; thick lines represent average perpendicular hand motion profiles. Similar results observed at T2. *Right panel*: Perpendicular hand deviations during loaded reaching movements to T3. Data were normalized by subtracting the average hand path during baseline movements from individual reaching trials with interaction loads. The robust controller's hand paths are less disturbed than the LQG controller when adaptation is incomplete (0, 25, 50, 75% adaptation). The robust and LQG controllers displayed similar hand path deviations when adaptation was complete (100% adaptation). Simulations produced qualitatively similar results at T2.

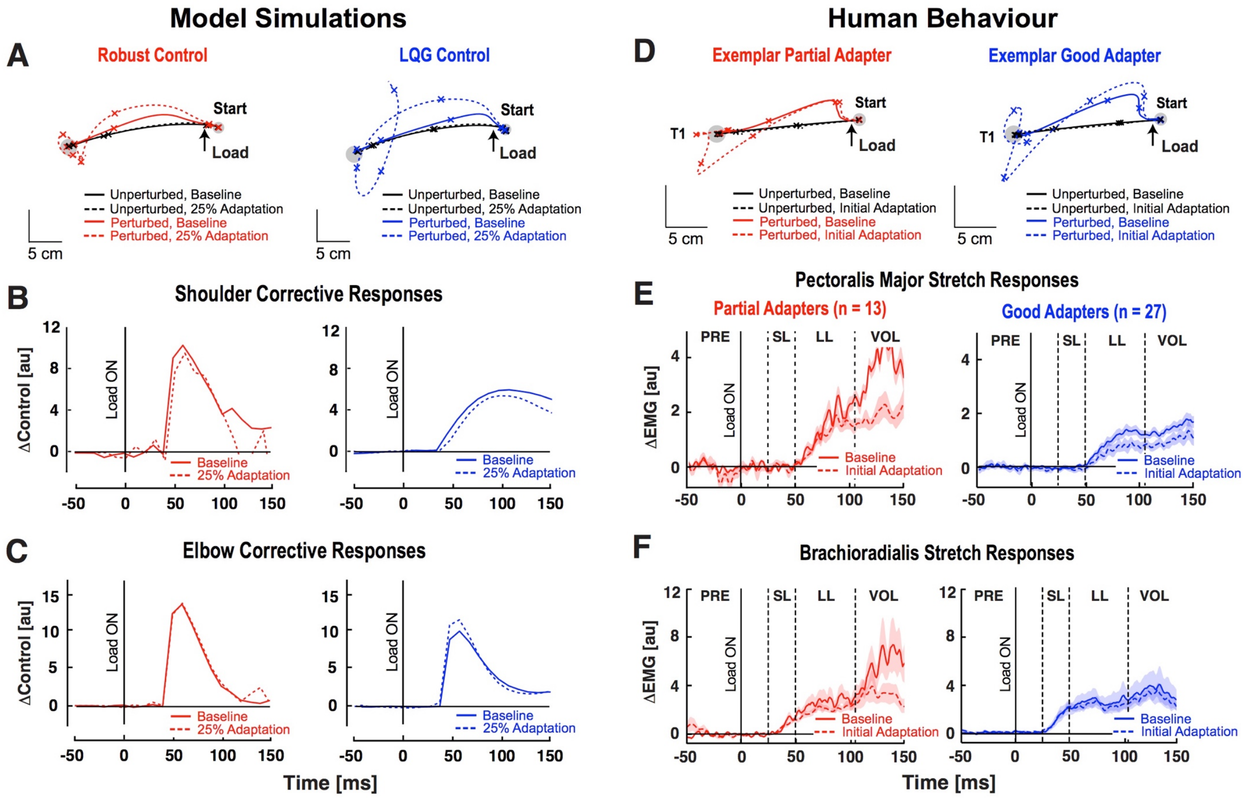


**Supplementary Fig. 6. Corrective responses to step-torque perturbations when initially exposed to novel interaction dynamics.** ***A***. Hand paths generated by robust control and stochastic (LQG) optimal feedback controllers during step-torque perturbation trials. Tick marks show hand positions every 200 ms. ***B.*** Average shoulder control responses to step-torque perturbations (n = 20). Data obtained by subtracting the average control responses in unloaded trials from individual step-torque perturbation trials (ΔControl). ***C.*** Elbow control responses during step-torque perturbation trials. Data are plotted in the same format as ***B***. ***D***. Hand motion of exemplar good and partial adapters during step-torque perturbation trials. Tick marks show hand positions every 200 ms. ***E.*** Pectoralis major stretch responses (mean ± SEM). Dashed vertical lines separate time windows of the muscle stretch response. ***F.*** Brachioradialis (BR) stretch responses. Data are plotted in the same format as ***E.***

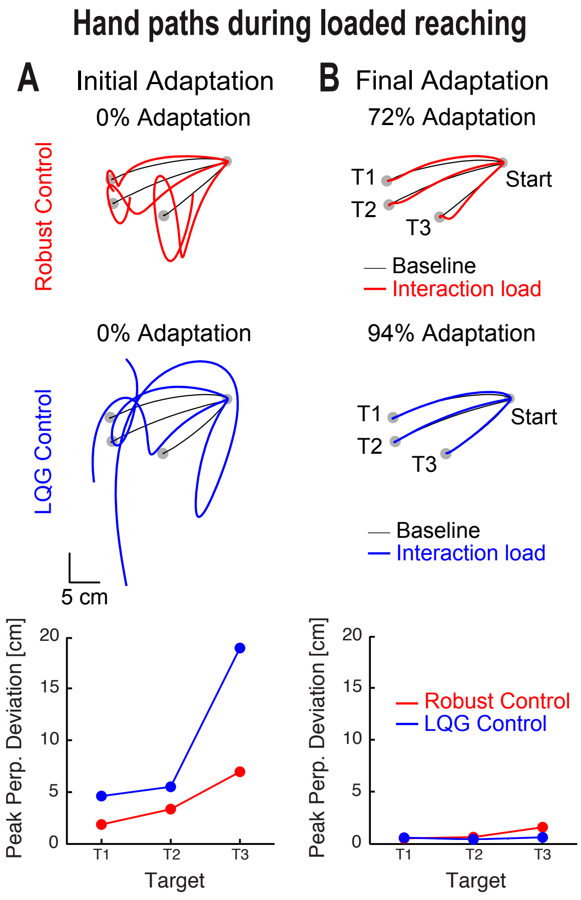


**Supplementary Fig. 7. Hand paths generated by robust and stochastic (LQG) optimal feedback controllers with different levels of knowledge of the novel interaction load. *A*.** Simulated hand paths during baseline and loaded reaching movements when the models had no knowledge of the interaction load (0% Adaptation). Thin black trace is the mean hand path during unloaded movements (n = 20 trials); thick coloured trace is the mean hand path while moving with novel interaction loads (n = 20 trials). Bottom panel shows average peak hand deviations during initial adaptation (0% Adaptation). ***B.*** Simulated hand paths when the models were constrained to the adaptation displayed by partial adapters (Robust-like control) and good adapters (LQG-like control). Data are plotted in the same format as ***A***.

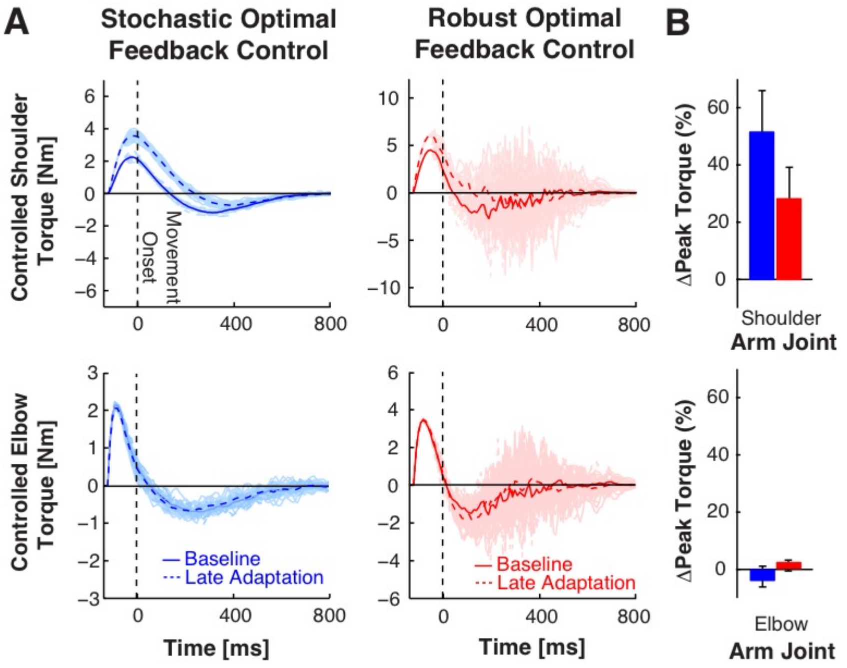


**Supplementary Fig. 8. Control torques generated by stochastic and robust optimal feedback controllers. A**. Shoulder (top row) and elbow control torques (bottom row) generated by stochastic and robust optimal feedback controllers while reaching to target 3 (T3). Thin lines represent individual trials (n = 20) in baseline and late adaptation. Thick solid lines represent average control torques generated during unloaded movements (i.e., baseline). Thick dashed lines represent the average control torques when the controllers were given adapted internal models. **B**. ∆Peak torques generated by stochastic and robust optimal feedback controllers between baseline and late adaptation. The accuracy of each controller's internal model was constrained to the amount of adaptation expressed by partial (Robust-like control) and good adapters (LQG-like control).

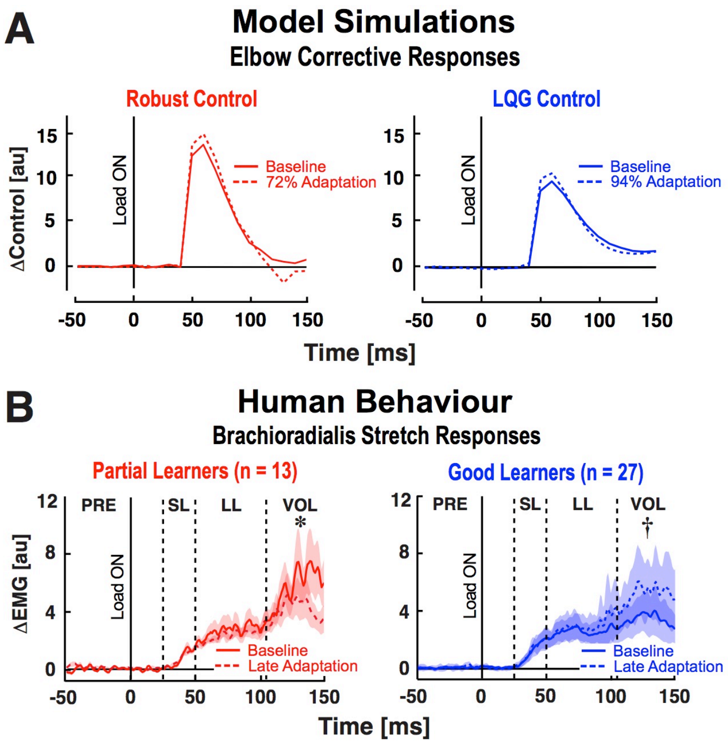


**Supplementary Fig. 9. Elbow muscle responses when disturbed by step-torque perturbations after adapting to novel interaction loads.** ***A.*** Elbow control responses generated by robust and stochastic (LQG) optimal feedback controllers during step-torque perturbation trials. Step-torque perturbations were simulated by abruptly changing the external torques acting on the shoulder and elbow joints during movement. Data obtained by subtracting the average control response during unperturbed reaching movements from individual step-torque perturbation trials. ***B.*** Brachioradialis (BR) stretch responses in human participants (mean ± SEM). Data are aligned to perturbation onset (solid vertical line, t = 0 ms). Dashed vertical lines separate different time windows of the muscle stretch response. $^{\dagger}p < 0.10$, $^{*}p < 0.05$.

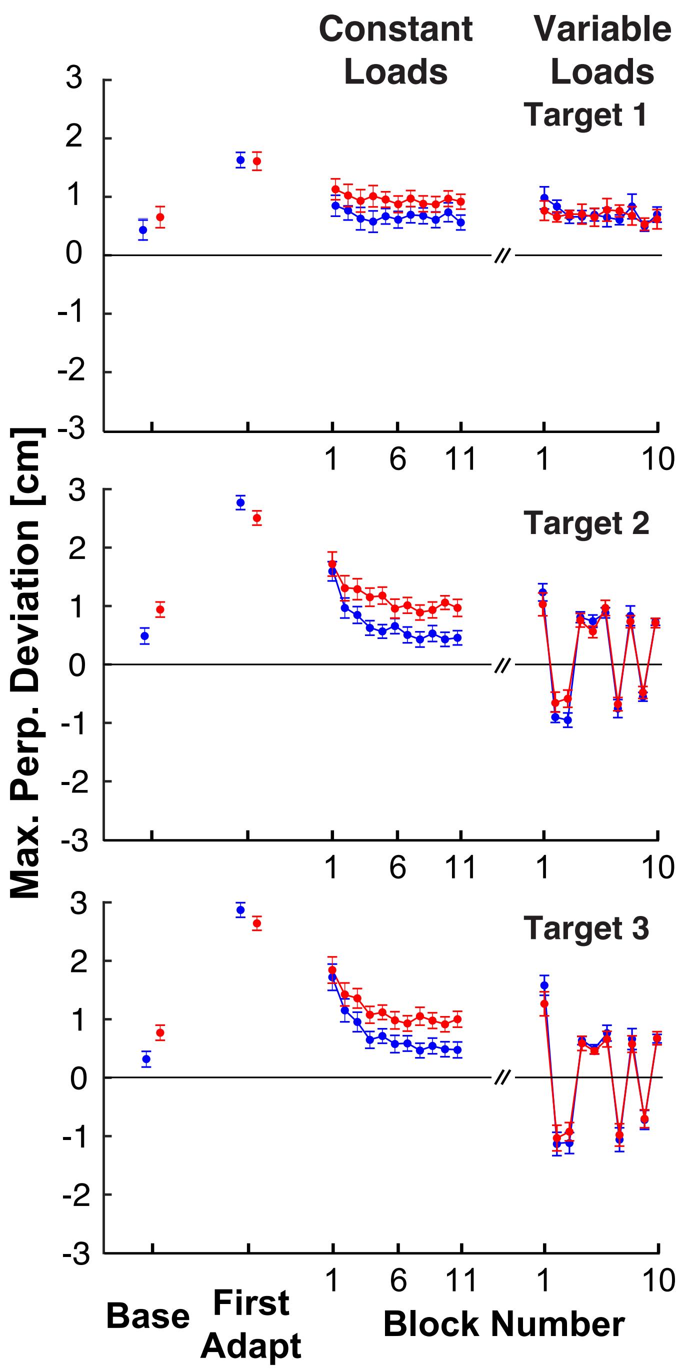


**Supplementary Fig. 10. Adaptation profiles in Experiment 2.** Maximum perpendicular deviations were calculated across late baseline, initial exposure to the interaction load at each target, across each block of trials during the adaptation phase and when exposed to rapidly varying interaction dynamics. Error bars represent ± 1 SEM.

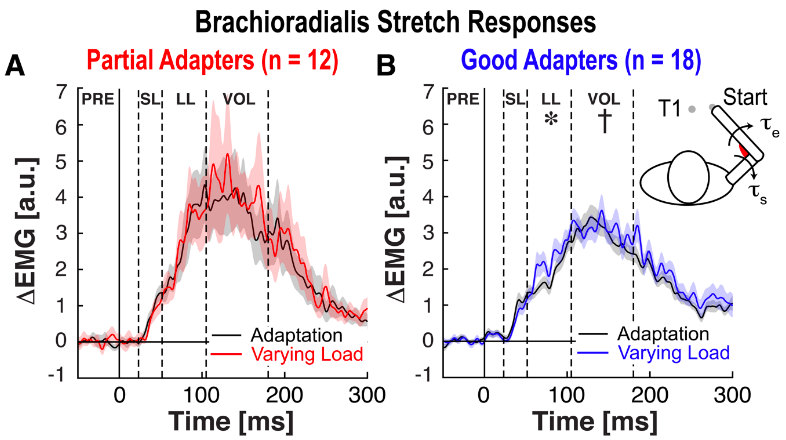


**Supplementary Fig. 11. Brachioradialis responses to step-torque perturbations when exposed to constant and varying interaction dynamics. A.** Brachioradialis response of partial adapters when step-torque perturbations extended the elbow while exposed to constant versus varying interaction dynamics. Dashed vertical lines separate different phases of the stretch response. **B.** Brachioradialis stretch responses when good adapters were exposed to constant versus varying interaction dynamics. Data are plotted in the same format as **A**. $^{*}p < 0.05$, $^{\dagger}p < 0.10$.
